# Circatidal control of gene expression in the deep-sea hot vent shrimp *Rimicaris leurokolos*

**DOI:** 10.1101/2024.01.12.575359

**Authors:** Hongyin Zhang, Takuya Yahagi, Norio Miyamoto, Chong Chen, Qingqiu Jiang, Pei-Yuan Qian, Jin Sun

## Abstract

Biological clocks are a ubiquitous feature of all life, enabling the use of natural environmental cycles to track time. Although studies on circadian rhythms have contributed greatly to the knowledge of chronobiology, biological rhythms in dark biospheres such as the deep sea remain poorly understood. Here, based on a laboratory free-running experiment, we reveal potentially endogenous rhythms in gene expression of the deep-sea hydrothermal vent shrimp *Rimicaris leurokolos*. Oscillations with ∼12-hour periods, likely reflecting tidal influence, greatly prevail over others in the temporal transcriptome, indicating *R. leurokolos* likely depends on a circatidal clock consisting of at least some components independent of the circadian clocks. The tidal transcripts exhibit an antiphased expression pattern divided into two internally synchronized clusters, correlated with wide-ranging biological processes that occur in the nucleus and cytoplasm, respectively. In addition, comparing the tidal transcripts with the ∼12-hour ultradian rhythms genes in fruit flies and mice shows large similarity, indicating the likely scenario of broad impact of the tide on the ∼12-hour oscillations across the metazoan. These findings not only provide new insights into the temporal adaptations in deep-sea organisms but also highlight deep-sea hydrothermal vent organisms as intriguing models for chronobiological, particularly 12-hour ultradian rhythms, studies.

## 1. Introduction

As an adaptation to natural periodic environmental changes, biological clocks have been evolving in all kingdoms of life to keep track of time [1–3]. Numerous organisms exhibit endogenous rhythms dominated by circadian periodicity of ∼24 hours at behavioral, physiological, and molecular levels, which help them anticipate daily changes such as light, temperature, and food availability [4–6]. To achieve this, animals rely on their internal circadian clock systems like transcription-translation feedback loops (TTFLs) that can be entrained to match the environmental cycles [7]. In dark biospheres such as the underground and deep-sea habitats, however, the absence of sunlight veils organisms there from the direct effects of everlasting day-night transitions. How organisms living in the dark schedule their activities through time, as an adaptation to darkness, remains obscure [8, 9].

The deep ocean, the largest but the least studied habitat on Earth [10], is of great interest for chronobiological research due to its distinct ecological environment [11]. With increasing depth, the sunlight intensity dramatically decreases and the visible light spectrum rapidly narrows to result in zero sunlight penetration below 1000 m even in the clearest oceanic waters [12]. From the ecological perspective, it is more appropriate for deep-sea organisms to synchronize their biological rhythms with other zeitgebers (e.g., tidal cycles) instead of the light cycle [13]. Indeed, various environmental fluctuations related to tidal cycles have been observed in the deep, especially in hydrothermal vent ecosystems [14–17]. Vent faunas rely strongly on their local environment, and survive on organic compounds produced by chemosynthetic microbes utilizing dissolved chemicals in the vent fluids [18]. Therefore, vent animals are likely to be affected by spatial and temporal variations in environmental factors modulated by the tide. Several previous studies have reported potential tide-related rhythms of behaviour and community dynamics in deep-sea organisms from chemosynthetic ecosystems [19–21], and a recent study revealed tidal cycles dominated the gene expression of the deep-sea vent mussel *Bathymodiolus azoricus* through periodic *in situ* sampling across 24.8 hours [22]. Although various evidence suggests that vent organisms exhibit biological rhythms in the wild, whether they need endogenous rhythms or clocks to accommodate and anticipate local environmental changes to maximize their fitness is still poorly understood.

In the Iheya North hydrothermal vent field of the Okinawa Trough (figure 1*a*), tidal modulation on the deep-sea current dynamics was observed previously [23]. According to a published long-term temperature measurement (Mean ± SD = 4.4 ± 0.1C) recorded at the periphery of the vent field where bathymodioline mussels typically occur, a strong semidiurnal tidal cycle (12.4-hour periodicity) was found in the environmental fluctuations as well (figure 1*b*). Unlike mussels at the vent periphery, however, the alvinocaridid shrimp *Rimicaris leurokolos* (formerly *Shinkaicaris leurokolos* [24]) lives in close proximity to the vigorously venting orifice (0.2–0.8 m and occasionally less than 0.2 m from the vent [25]) where the hot fluid and cold surrounding seawater mix spontaneously and dramatically [26] (figure 1*c*). In the *R. leurokolos* colony, environmental factors fluctuate frequently and dramatically at the seconds-minutes scale, which adds unpredictable noises to the tidal cycles. In fact, the temperature could rapidly fluctuate between 7.63 to 17.74°C (Mean ± SD = 10.16 ± 1.71°C) within approximately 2.5 minutes (figure 1*d*, electronic supplementary material, data S1). As such, *R. leurokolos* is likely unable to directly use the unsteady, highly variable environmental factors as zeitgebers to entrain its biological rhythm. However, patterns of venting activity across a longer time-span is known to be linked to tidal cycles [27]. Therefore, we hypothesised that an internal clock may integrate useful periodic cues whether they are clear (e.g., pressure) or noisy (e.g., water chemistry), and thus help hot vent shrimps adapt and keep track of periodic environmental changes.

**Figure 1.**
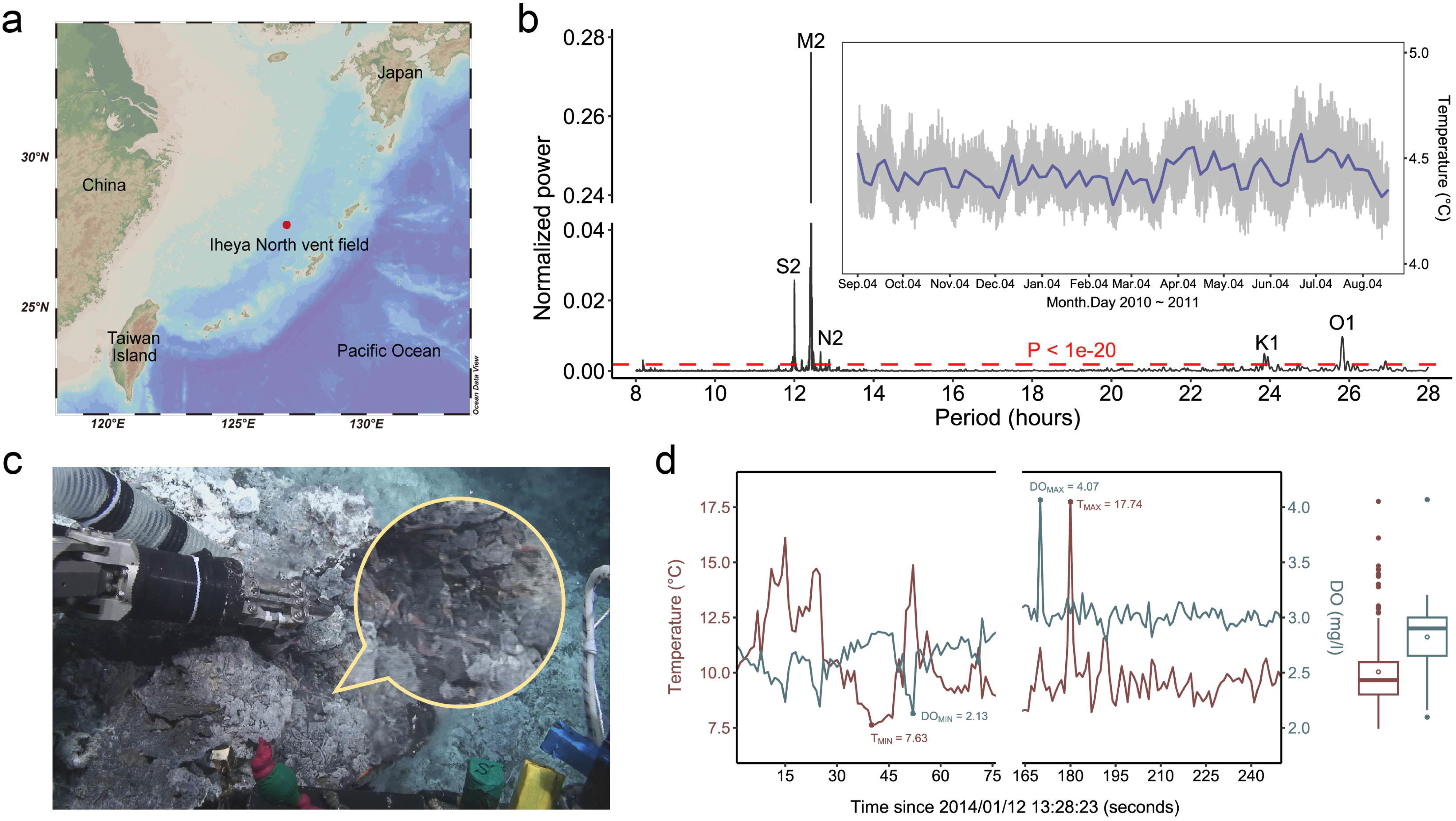
Deep-sea chronobiology study on the hydrothermal vent shrimp *Rimicaris leurokolos*. (*a*) This work was focused on the Iheya North hydrothermal vent field, Okinawa Trough. (*b*) Time series of published temperature data covering 351 days recorded at the peripheral area of the active NBC mound vent. Lomb-Scargle (LS) periodogram analysis reveals a strong semidiurnal tidal cycle dominated by the M2 tidal constituent. M2, principal lunar semidiurnal constituent; S2, principal solar semidiurnal constituent; N2, larger lunar elliptic semidiurnal constituent; K1, luni-solar declinational diurnal constituent; O1, Lunar declinational diurnal constituent. (*c*) *R. leurokolos* shrimps (in the yellow circle) live in a hot zone near vent orifice where hydrothermal fluids mix with cold surrounding seawater spontaneously. (*d*) Time series of *in situ* temperature and dissolved oxygen records during 162 seconds when the sensor was placed in the *R. leurokolos* colony showing frequent environmental fluctuations with high variability.

Although circadian patterns are dominant in terrestrial and coastal animals, recent studies revealed robust ultradian rhythms (e.g., ∼12-hour cycling gene expression) in model organisms such as mice and fruit flies [28, 29]. These rhythms were found to be cell-autonomous, driven by a distinct 12-hour pacemaker largely independent from the circadian clock. As many homologous genes were found rhythmic with a period of ∼12 hours both in terrestrial and marine animals, a hypothesis was then proposed that the circatidal rhythms in marine animals and ∼12-hour rhythms in terrestrial animals are evolutionarily conserved [30]. However, given the dominant effects of the ∼24-hour circadian system, it is challenging to distinguish the ultradian rhythms from circadian rhythms in terrestrial or coastal animals. The hot vent shrimp studied herein lives in the deep sea environment lacking day-night cycles, and thus provides an opportunity to examine this hypothesis.

Here, through a laboratory free-running experiment across 72 hours, we addressed the biological rhythms of gene expression in the vent shrimp *R. leurokolos* and tested if it can maintain rhythmicity under constant conditions. Our work reveals free-running rhythms dominated by 12-hour tidal patterns in the transcriptome of *R. leurokolos*, which are likely driven by an unidentified endogenous clock including but not limited to circatidal components. The tidal transcripts exhibit a bimodal gene expression pattern, and influence broad cellular processes such as DNA replication, DNA repair, regulation of transcription, transport, cell cycle, and responses to environmental stresses. In addition, we found *R. leurokolos* shares many common genes and biological processes in the tidal rhythms with the ∼12-hour rhythms in sea anemones, fruit flies, and mammals, suggesting the evolutionary conservation of ∼12-hour gene expression across a broad range of phyla. These findings highlight the prospect for deep-sea organisms in chronobiology, especially in the evolution and molecular mechanisms of ∼12-hour ultradian rhythms.

## 2. Materials and methods

### (a) Environmental temperature monitoring

The local fluctuations in the temperature of a *Rimicaris leurokolos* colony on the NBC chimney (126°53.80’E, 27°47.45’N; 982 m deep) of the Original Site, Iheya North vent field were measured using a RINKO dissolved oxygen and temperature sensor (JFE Advantech Co., Ltd.) during dive #1610 of the remotely operated vehicle (ROV) *Hyper-Dolphin* on-board R/V *KAIYO* cruise KY14-01 for 162 seconds, and data were recorded every second (electronic supplementary material, data S1). Published temperature data in a peripheral area of the original site, Iheya North vent field far away from active chimneys, which were recorded continuously every 10 min from the 4th of September 2010 to the 20th of August 2011, were re-analyzed [31]. We used the Lomb-Scargle periodogram to detect periodicity in the temperature time series.

### (b) Sampling procedure and experiment design

A total of 184 adult individuals of *Rimicaris leurokolos* were collected by a suction sampler from the top of the NBC chimney at the Original Site, Iheya North hydrothermal field during dive #1613 of ROV *Hyper-Dolphin* on-board R/V *KAIYO* in the cruise KY14-01. Specimens of *R. leurokolos* were kept in canister bottles on the ROV prior to recovery on-board the research vessel. After sampling, it took about 2 hours (15:17–17:07 JST) for the ROV to be recovered. Once on-board the ship, under natural light, 154 live shrimps, excluding 25 dead individuals and 5 ovigerous females, were immediately transferred into a lightproof seawater container. This container was placed in a dark cold room at 4°C by 19:30 JST for the subsequent free-running experiment.

The free-running experiment began at 22:00 JST, aiming to investigate the gene expression of *R. leurokolos* in a constant environment over time. During the entire experiment light was turned off, although a faint orange light from the power strip remained lit. As this was very dim and with a wavelength close to red light, the impact of this should be minimal. At each sampling timepoint, shrimps were collected by a meshed scoop, dissected with scissors without anesthesia into cephalothorax and abdomen, and fixed every 4 hours for a total duration of 72 hours. The cephalothorax of each individual were preserved in RNAlater (Invitrogen) and used for RNA-Seq in this study. During the sampling, we did not provide any food source as the previous laboratory study on behaviours of vent shrimps revealed that the introduction of food largely increased the activities of these shrimps [32]. As a possible zeitgeber [33], an external food source may give an opportunity for shrimps to synchronize their activities and even gene expression at a certain time, which should be avoided in the free-running experiment. Despite that, *R. leurokolos* may feed on bacteria attached to other individuals. We did not find any dead shrimp during the free-running experiment, which might reflect the health conditions of *R. leurokolos*.

### (c) RNA extraction, sequencing and assembly

Total RNA was extracted from individuals using TRIzol reagent (ThermoFisher) following the manufacturer’s protocol. The mRNA from each shrimp cephalothorax was enriched by Oligo-dT probes and further reverse transcribed to first-strand cDNA for library construction. All cDNA libraries were sequenced on an Illumina NovaSeq 6000 platform with 150 bp paired-end mode. A total of 57 RNA-Seq samples were generated from *R. leurokolos* during 72 hours of sampling at 4-hour intervals with three replicates per time point. Raw reads were firstly trimmed using Trimmomatic version 0.39 [34], and then potentially associated bacteria contamination reads were removed with Kraken2 version 2.0.8-beta [35]. *De novo* assembly was performed with Trinity version 2.11.0 [36] (set jaccard clip option). Salmon version 1.4.0 [37] was used to quantify the expression of each assembled transcript.

### (d) Gene quantification and annotation

The TMM cross-sample normalization method [38] was applied to remove the sequencing variations and potential “batch effects” among samples using the abundance_estimates_to_matrix.pl script in the Trinity toolkit. In addition, lowly expressed transcripts with 0 counts in at least 38 samples were filtered out, leaving 23,905 transcripts for subsequent analyses. The predicted proteins were annotated by searching the candidate protein sequences against the NCBI non-redundant (NR) database using BLASTP with DIAMOND version 2.0.13 [39] with ultra-sensitive mode and E-value threshold of 1 × 10^-5^, the Kyoto Encyclopedia of Genes and Genomes (KEGG) database via KEGG Automatic Annotation Server (KAAS), and the Gene Ontology (GO) via BLAST2GO version 6.0 [40]. The gene enrichment analyses were performed with ClusterProfiler version 4.2.2 [41].

### (e) Gene expression and detrend analyses

Principle components analysis (PCA) was performed using the plotPCA function in R package DESeq2 version 1.34.0 [42]. Permutational multivariate analysis of variance (PERMANOVA) and analysis of similarities (ANOSIM) were applied to test the statistical significance of the PCA results. The potential monotonic trends in the raw gene expression of all transcripts were determined using the Mann-Kendall trend test, and we pinpointed two overall clusters of transcripts that had either an upward trend (S score > 0 and *P*-value < 0.05) or a downward trend (S score < 0 and *P*-value < 0.05).

To remove the trend and produce a steady baseline for subsequent rhythm analyses, we used CEEMDAN [43], an adaptive decomposition method designed for non-stationary and non-linear time series signals, which minimizes the impact of distorting the period, phase or amplitude change of the oscillations. By this method, each gene expression data could be decomposed into three intrinsic mode functions (IMFs), and a residual that represents the trend (electronic supplementary material, figure S1). In addition to the potential periodic patterns, the three IMFs are also likely to contain noises. We then selected the relevant IMFs and reconstructed the detrended gene expression data under the criterion suggested by Ayenu-Prah & Attoh-Okine [44]. To verify the effectiveness of CEEMDAN detrend analysis, we compared the rhythm detection results using gene expression data before and after detrending. Rhythmic datasets identified using raw and detrended data share many common transcripts, and we detected more rhythmic transcripts through the detrended data (electronic supplementary material, figure S2). In addition, using the detrended gene expression, we detected extra periodic patterns in some transcripts that had a significant monotonic trend within the raw gene expression, which were hard to recognize using the raw data (electronic supplementary material, figure S3). These findings suggest that the detrend analysis increases the power of rhythmicity detection methods and thus helps us to get a clearer picture of the periodic patterns within the gene expression data. The ceemdan detrend was performed with the R package Rlibeemd version 1.4.2 [45] (set parameters: num_imfs = 0; noise_strength = 0.4; S_number = 0; num_siftings = 50; ensemble_size = 250)

### (f) Rhythm detection analyses

For *R. leurokolos*, we knew nothing about its biological rhythm, and no golden-standard circadian or non-circadian genes were available for evaluating the detection power (such as AUC values and ROC curves) of different algorithms [46, 47]. Thus, we assessed various rhythm detection methods based on our experiment design (for details see electronic supplementary material, table S1). Overall, we chose four rhythm detection algorithms in this study, the mathematic method eigenvalue/pencil [28] (carried out with customized scripts), the Extended Circadian Harmonic Oscillator application version 4.0.1 [48] (ECHO), and two non-parametric statistical methods RAIN version 1.28.0 [49] and empirical JTK_CYCLE version 1.0 [50] (eJTK), which were designed to detect asymmetric waveforms. For eigenvalue/pencil and ECHO, rhythmic transcripts with the periodicity of 10-14 hours and 22-26 hours were considered as tidal and daily oscillated [51]. Oscillations with the largest amplitude in one transcript for eigenvalue/pencil were considered the dominant oscillation [52]. For RAIN and eJTK, rhythmic transcripts with a period of 12 hours and 24 hours were considered tidal and daily cycling. Multiple-testing was carried out to adjust *P*-values using the Benjamini-Hochberg method for RAIN and ECHO, and an empirical method based on the gamma distribution for combining dependent *P*-values for eJTK. Transcripts with an FDR ≤ 0.05 were considered statistically rhythmic. The root mean square (RMS) amplitude and Rhythmicity Index (RI) of rhythmic transcripts were calculated by LimoRhyde2 version 0.1.0 [53] and autocorrelation analysis, respectively. We also used ECHO to fit data with an extended fixed amplitude model and categorize the oscillation type using amplitude change coefficient (γ) as forced (−0.15 < γ < −0.03; amplified amplitude), harmonic (−0.03 ≤ γ ≤ 0.03; steady amplitude), damped (0.03 < γ < 0.15; reduced amplitude), overexpressed (γ ≤ −0.15), and repressed (γ > 0.15), of which the latter two are considered arrhythmic [48]. The results of rhythm detection analyses are listed in electronic supplementary material, data S2.

### (g) Identification of putative circadian clock genes

We searched for putative circadian clock genes in our *de novo* assembled transcriptome using the protein Basic Local Alignment Search Tool (BLASTP) carried out by DIAMOND with a series of known core clock and clock-associated genes as queries (electronic supplementary material, data S3). For candidate circadian clock transcripts, sequences were scanned against the coding regions and conserved functional domains using the InterProScan tools [54]. To determine the sequence homology of candidate transcripts, we further conducted the phylogenetic analysis. The amino acid sequences were first aligned using MAFFT version 7.490 [55] and then trimmed using trimAl version 1.4.1 with the ‘-automated1’ option [56]. The maximum-likelihood trees were then constructed using the IQ-Tree version 2.1.3 [57] with the ultrafast bootstrap approximation (UFBoot) and ModelFinder for examining the best-fit model. No *Bmal*/*Cycle* transcript was identified in *R. leurokolos* and the *Clock* transcript exhibited extremely low expression that rendered it unfeasible for rhythm analysis. Overall, we reconstructed four phylogenetic trees using the Q.insect+Γ4 model for the Clock and Bmal/Cycle tree, the Q.insect+I+Γ4 model for the Period tree and the Timeless/Timeout tree, plus the LG+R4 model for the Photolyase/Cryptochrome tree. In the identified circadian clock genes, the *Clock* transcript exhibited extremely low expression and was thus excluded from the rhythm detection. Visualization of all phylogenetic trees was performed with ggtree version 3.8.2 [58].

### (h) Conserved rhythmic orthologs analysis

The 12-hour rhythmic gene expression data of mice liver and *Drosophila* S2 cell used in this study were published previously [28, 29]. Together with the *de novo* transcriptome of *R. leurokolos* assembled in this study, the protein sequences from mouse genome assembly GCF_000001635 and fruit fly genome assembly GCF_000001215 were used to construct orthogroups using Orthofinder version 2.5.4 [59] with default settings. The conserved rhythmic analysis was performed with the single-copy orthologous genes. Pathway network enrichment analysis of conserved single-copy orthologous genes was performed with aPEAR version 1.0.0 [60].

## 3. Results and discussion

### (a) Transcriptome analysis of deep-sea vent shrimp *R. leurokolos*

To study the potential endogenous rhythms in *R. leurokolos*, we carried out a 72-hour free-running experiment with 4-hour sampling intervals under constant conditions. According to previous studies on crustaceans, we worked on the whole cephalothorax that contained the nervous and visual system where the central clock likely lies, since the exact location in crustaceans remains unclear [61]. The reference *de novo* transcriptome contained 90,755 predicted open reading frames (ORFs), which showed a 92% complete BUSCO score. Candidate protein-coding transcripts with low expression were filtered out, leaving 23,905 transcripts with a 60.4% annotation rate for rhythmic analyses. A principal components analysis (PCA) of normalized raw gene expression showed a rough trend from the start of sampling (0 hours) to the end (72 hours, electronic supplementary material, figure S4). Performing the Mann-Kendall trend test on the raw dataset, we found that 10,273 transcripts (i.e., 42.97% of the transcriptome) had a significant monotonic trend (*P*-value < 0.05). The trend was then removed using CEEMDAN [43] to produce a steady baseline and enhance the strength of rhythm detection methods (electronic supplementary material, figure S5). In the PCA results plotted using detrended gene expression, the trend disappeared and sample points from different sampling times clustered together (figure 2*a*), indicating the shrimps were seemingly experiencing the same conditions.

**Figure 2.**
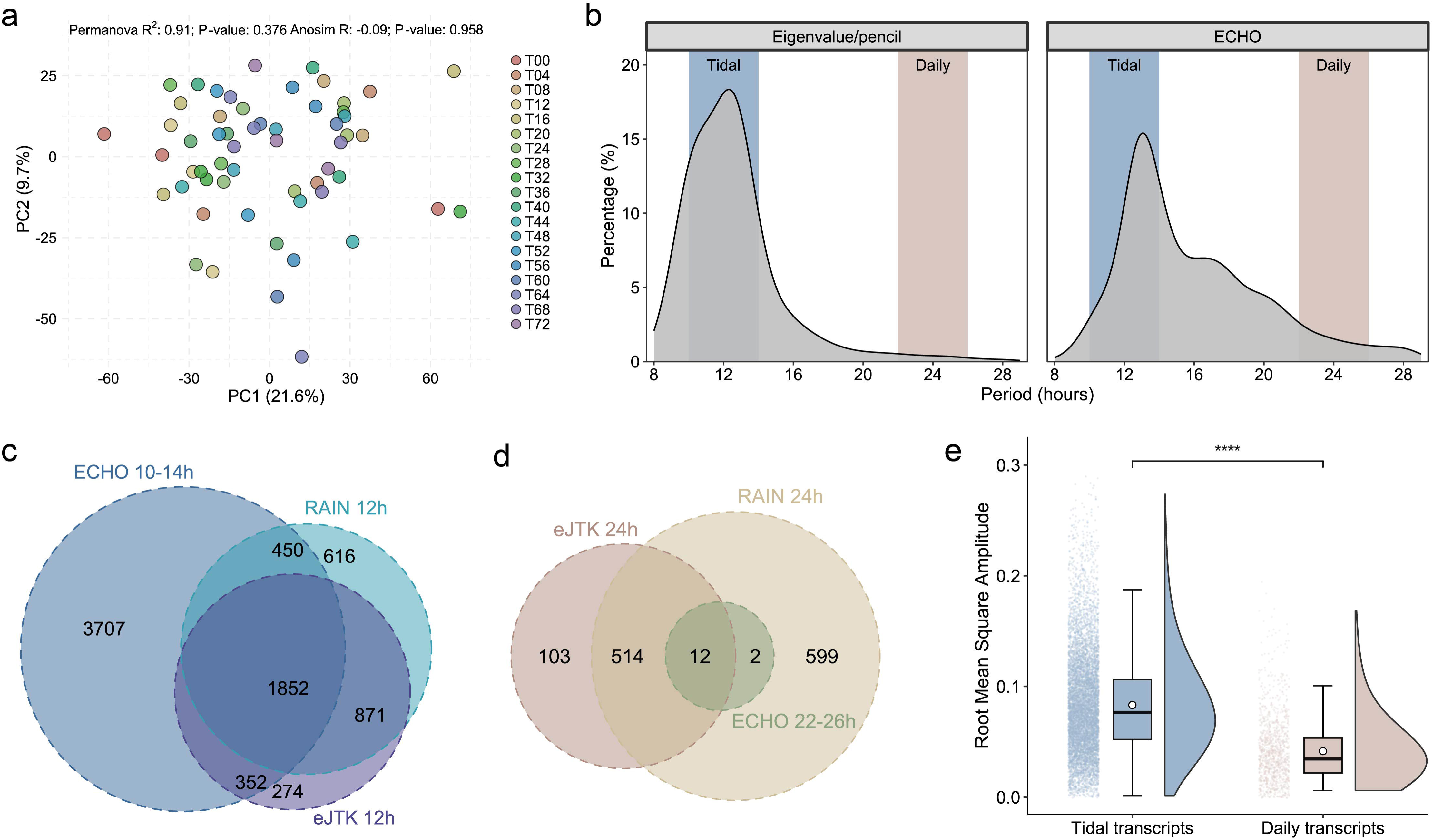
The *Rimicaris leurokolos* temporal transcriptome was dominated by tidal oscillations under constant conditions. (*a*) Principal components analysis (PCA) plot showing the detrended transcriptomic data from the free-running experiment samples (n = 57). Significance of inter-group differences was examined by the PERMANOVA (*P*-value = 0.376) and ANOSIM (*P*-value = 0.958) analysis both on Bray-Curtis dissimilarities. (*b*) Distribution of the periods of oscillations identified by the eigenvalue/pencil method and the ECHO application. Oscillations with a period range of 10-14 hours and 22-26 hours are regarded as tidal and daily cycling, respectively. (*c,d*) Venn diagrams detailing the number of rhythmic transcripts (FDR < 0.05) identified by ECHO, RAIN, and eJTK in the (*c*) tidal range and (*d*) daily range. (*e*) Raincloud plot of the root mean square (RMS) amplitude for 12-hour tidal transcripts and 24-hour daily transcripts. The stars indicated the level of significance: **** *P*-value < 0.0001.

### (b) General biological rhythms in the temporal transcriptome of *R. leurokolos*

We first used the eigenvalue/pencil [28] and ECHO [48] algorithms to carry out unbiased identification of any potential periodic oscillations in the gene expression of *R. leurokolos*. The results uncovered similar rhythmicity that most cycling transcripts were identified as ultradian rhythms, which oscillated in the tidal range of 10-14 hours, while much fewer cycling transcripts oscillated in the daily range of 22-26 hours (figure 2*b*). The eigenvalue/pencil algorithm was able to identify superimposed oscillations from one cycling transcript. For cycling transcripts with multiple components composed of tidal and daily oscillations, the amplitude of the tidal component is significantly larger, suggesting stronger rhythmicity (electronic supplementary material, figure S6). Despite the high sensitivity of detecting weak ultradian rhythms, the eigenvalue/pencil method could have high positive false rates [52]. To validate the results, we then detected cycling transcripts with two robust nonparametric algorithms, RAIN [49] and empirical JTK_CYCLE [50] (eJTK). Searching for cycling transcripts (FDR < 0.05) with periods in the range of 8-28 hours at regular intervals of 4 hours, we found the number of rhythmic transcripts identified by RAIN and eJTK both reached a peak at the 12-hour period that corresponded to a tidal cycle (electronic supplementary material, figure S7). Interestingly, ECHO shared many common cycling transcripts with RAIN (60.8% of transcripts) and eJTK (65.8% of transcripts) but detected numerous extra tidal transcripts, likely due to the inclusiveness of identifying different harmonics of oscillating elements [62]. Merging transcripts that were identified as rhythmic by at least one method, we identified a total of 8,122 rhythmic transcripts (33.98% of the transcriptome) in the tidal range (figure 2*c*), versus 1,230 rhythmic transcripts (5.15% of the transcriptome) in the daily range (figure 2*d*). Then we calculated and compared the root mean square (RMS) amplitude between tidal and daily transcripts, and found the tidal transcripts had a significantly larger amplitude than daily ones, indicating tidal cycles contributed more biologically relevant effects (figure 2*e*). The list of rhythmic transcripts is provided in electronic supplementary material, data S2.

These results demonstrate sustainable endogenous rhythmic gene expression in *R. leurokolos* under constant conditions. The temporal transcriptome is dominated by the approximately 12-hour tidal rhythms, which greatly prevail over other oscillations with different periods, especially the 24-hour daily transcripts that are likely unrelated to the circadian rhythms.

### (c) Patterns of tidal gene expression in *R. leurokolos*

To uncover the biological rhythms ticking in the deep, we then focused on the dominant tidal rhythms in the temporal transcriptome of *R. leurokolos*. Our 72-hour duration of sampling covered approximately six tidal cycles, allowing us to test the stability and robustness of rhythmicity in the tidal gene expression [63]. To capture the amplitude changes over time, we fitted the gene expression to an extended fixed-amplitude model using ECHO [48] and categorized the results into five types of oscillation groups (details see Materials and methods). Near half of the tidal transcripts (n = 3,294) were identified as damped type, showing a reduced amplitude (figure 3*a*).

**Figure 3.**
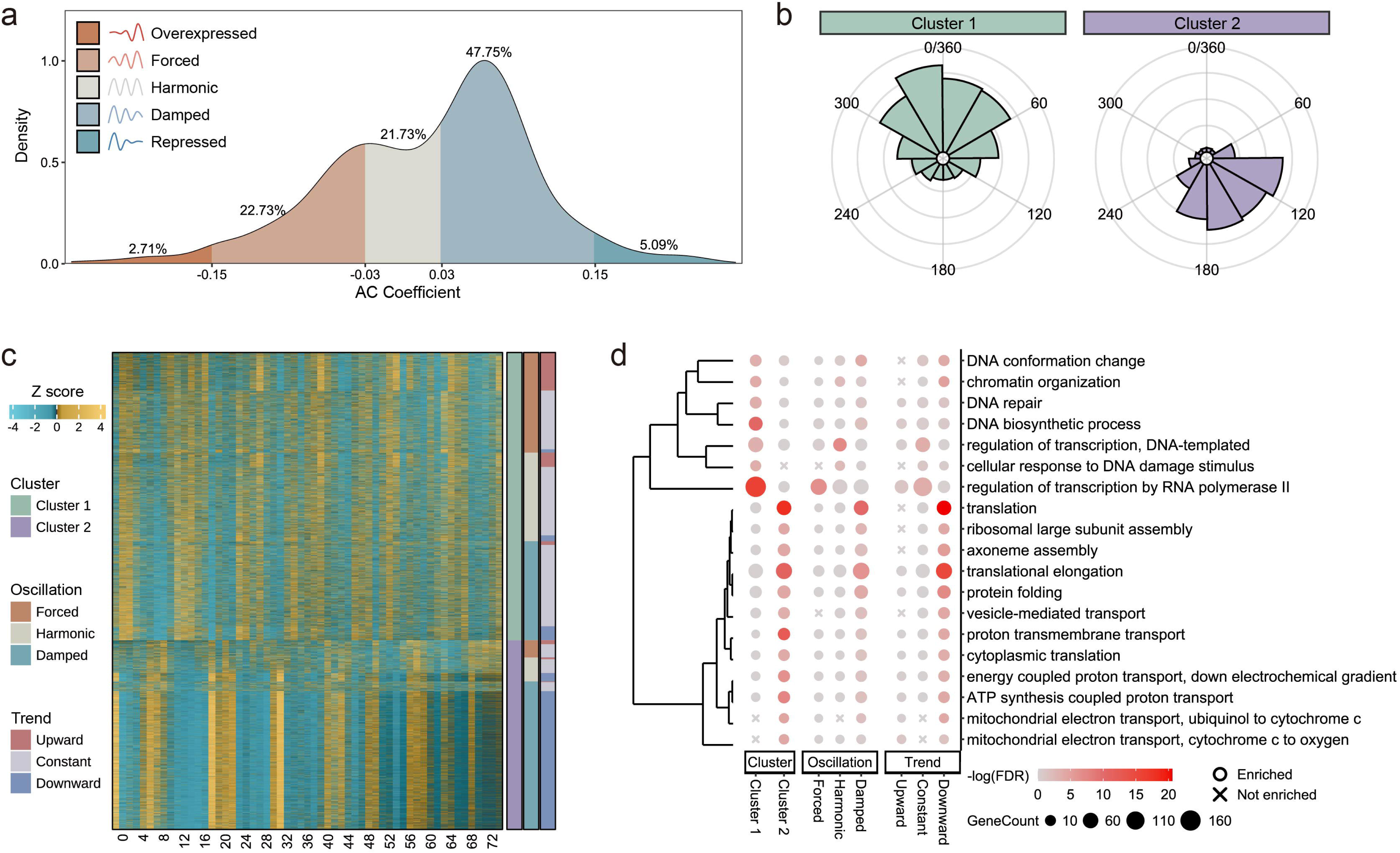
Free-running tidal transcriptomic patterns of *Rimicaris leurokolos*. (*a*) Density distribution based on amplitude change coefficient (γ) of tidal transcripts detected by ECHO. Oscillation types are categorized as forced (−0.15 < γ < −0.03; amplified amplitude), harmonic (−0.03 ≤ γ ≤ 0.03; steady amplitude), damped (0.03 < γ < 0.15; reduced amplitude), overexpressed (γ ≤ −0.15), and repressed (γ > 0.15), and the latter two are considered arrhythmic. (*b*) Rose chart showing phase distribution comparison of tidal transcripts in cluster 1 and cluster 2. For each oscillation, the phase is divided by period length and then times 360 (degrees) to normalize phases of transcripts with different period lengths. (*c*) Heatmap showing highly synchronized gene expression of tidal transcripts (ECHO FDR < 0.05). Each row represents a transcript with three individuals per time point. Transcripts are clustered and ordered by phases, oscillation type, and monotonic trend. The expression of each gene is Z-score normalized. (*d*) Bubble diagram showing significantly enriched gene ontology (GO) terms related to biological processes of tidal subgroups categorized by three classification schemes (from left to right: cluster, oscillation type, trend). Cross symbols represent a lack of enriched terms in the corresponding subgroup.

Although there was a significantly amplified or reduced amplitude in at least 75% of tidal rhythmic transcripts, the period and phase of most tidal cycles were seemingly fixed. We divided the tidal transcriptome into two clusters by gene expression using K-Means clustering (electronic supplementary material, figures S8*a* and S8*b*). The two clusters showed a largely antithetical gene expression pattern over time, and transcripts within each cluster were peaking at the same time, resulting in an almost complementary phase distribution in one tidal cycle for the two clusters (figures 3*b*). This bimodal expression pattern is widely observed in the gene expression of endogenous rhythms. For example, terrestrial organisms such as mice and baboons experience transcriptional ‘rush hours’ preceding dawn and dusk, which is considered to assist in preparing for the impending change in lighting [5, 6]. In the hydrothermal vent, tidal cycles could strongly affect variations in environmental factors such as temperature, water chemistry and dissolved oxygen. Vent fauna would thus experience two opposite environmental conditions in a tidal cycle that one is high temperature, high sulfide, low oxygen, etc., and another is comparatively low temperature, low sulfide, high oxygen, etc. In this case, the bimodal tidal expression pattern could help *R. leurokolos* save energy during the transition periods between acrophase and trough in the tidal cycle, and make preparations for the upcoming significant environmental changes [51, 64, 65].

The rhythmicity of 3,837 transcripts in cluster 1 was relatively stable with almost equivalent amounts of modest forced, harmonic, and damped oscillations, whereas the 2,524 transcripts in cluster 2 were nearly occupied by damped oscillations that decreased obviously in amplitude, particularly during the second half of experiment (figure 3*c*). Indeed, there was seemingly a strong correlation between trend directions and oscillation types (electronic supplementary material, figure S8*c*). In other words, a tidal transcript with a decreasing expression trend was more likely to have a reduced amplitude, and vice versa. This likelihood was extremely high in cluster 2 where over 94% of tidal transcripts with a downward trend were also damped. Since we isolated *R. leurokolos* shrimps from the hydrothermal vent and cultivated them in the lab under constant conditions for 72 hours, there were unsurprisingly considerable changes in the gene expression over time corresponding to transcriptional responses to potential stresses such as decompression. Therefore, the amplitude likely decreases as a byproduct of declining gene expression levels, especially for most transcripts in cluster 2. Although desynchronization between different tissues of *R. leurokolos* could lead to arrhythmic gene expression in our free-running study, we still detected a considerable number of rhythmic transcripts with highly synchronized expression, suggesting the effect of desynchronization was likely minor.

### (d) Wide range of biological processes related to tidal cycles in *R. leurokolos*

The tidal rhythmic transcripts could be categorized by gene expression, oscillation types, and original trends, all of which included several subgroups that likely reflected similar or different biological relevance. To search for potential biological functions modulated by tidal rhythms, we performed functional enrichment analyses over the tidal transcriptome based on these three classification schemes using clusterProfiler [41]. Gene ontology (GO) and KEGG pathway analyses revealed that tidal rhythms generally affected various biological processes such as DNA repair, regulation of transcription, translation, transport, organelle organization, and citrate cycle (figure 3*d*, electronic supplementary material, figure S9). There was an expected high correlation between the biological functions of classified subgroups. For example, terms enriched in cluster 2 were more or less the same as those in the damped or downward subgroup. Notably, rhythmic processes related to the nucleus such as chromatin organization, DNA biosynthetic process, and transcription were uniquely enriched in cluster 1; whereas rhythmic processes supported by the cytoplasm such as translation, protein folding, vesicle-mediated transport, and oxidative phosphorylation were highly enriched in cluster 2. Thus, it is likely that *R. leurokolos* shrimps can organize various rhythmic biological processes in terms of spatial and temporal expression patterns to coordinate their internal physiology, regardless of the presence of clear zeitgebers.

In hydrothermal vents, particularly near the fluid orifice where *R. leurokolos* inhabits, hot, anoxic fluids with abundant potentially toxic chemical substances (e.g., hydrogen sulfide) mix dramatically with the cold surrounding seawater, resulting in an extreme and variable environment. Vent organisms have evolved various adaptive strategies at morphological, physiological, and biochemical levels [18]. We hypothesized that intrinsic rhythms could help vent animals schedule their activities to cope with the harsh environment. As expected, we found many tidally oscillating transcripts were related to various cellular stress response processes such as sulfur metabolism, reactive oxygen species (ROS) homeostasis, cellular detoxification, heat and hypoxia response and innate immune response (electronic supplementary material, figures S10*a*-*d*). The representative tidal genes related to stress responses include *Sqor*, *Cat*, *Sod*, *Gst*, *Gpx*, *Egln1*, *Dnaja1*, *Hsp90*, *Ctsl*, among others. Most of these transcripts showed damped rhythmic expression over time under constant conditions, suggesting that *in situ* environmental cues were necessary for the maintenance and synchronization of rhythmic processes related to stress responses. The results reveal broad tidal regulation of gene expression in the cellular response to various environmental stresses, suggesting an adaptive advantage for *R. leurokolos* living in harsh, fluctuating environments.

Mounting evidence suggests the presence of cross-talking between circadian clocks and the cell cycle [66, 67]. Here, we found the annotation of many tidal transcripts was related to the cell cycle (electronic supplementary material, figures S11*a*-*c*). The transition between the G1 and S phase is controlled and regulated by key genes involved in the checkpoint pathways [67–69], of which several transcripts were tidally oscillating, including *Ccnd2*, *TP53*, *Mycbp2*, *Rbl1*, *E2f2*, *Tipin*, *Mdm2*, among others. In addition, genes that encoded major components of the RAD9-RAD1-HUS1 clamp complex (e.g., *Rad9a*, *Rad1*, *Hus1*), which play pivotal roles in DNA repair, also oscillated in the tidal range [70]. As crucial elements of the spindle assembly checkpoint (SAC), genes such as *Mad2*, *Bub1*, and *Bub3* that regulate chromosome separation were also found to be tidally cycling [71]. Taken together, tidal rhythms appear to generally regulate the progression of the cell cycle in *R. leurokolos* to cope with DNA damage derived from periodic intrinsic and environmental stresses.

### (e) Possible molecular basis and zeitgebers of biological rhythms in *R. leurokolos*

We searched for and identified *R. leurokolos* homologs of the canonical circadian clock genes that form the molecular oscillator in the fruitfly *Drosophila melanogaster*, including *Clock (Clk)*, *Period (Per)*, *Timeless (Tim)*, *Cryptochrome 2* (*Cry2)*, *Clockwork orange* (*Cwo)*, *Doubletime (Dbt)*, *Vrille* (*Vri*), and *Pdp1*ε (electronic supplementary material, figures S12*a-d* and data S3). Indeed, all identified *R. leurokolos* circadian clock gene homologs lacked 24-hour rhythmicity (figure 4). This may provide intrinsic fitness for these hydrothermal vent shrimps that are likely completely isolated from the surface as adults [72]. Moreover, most clock genes showed a lack of rhythmicity in the tidal range as well. Although the transcriptional repressor *Period* and *Doubletime*, which encodes a kinase that phosphorylates the PER protein [73], were statistically oscillated in the tidal range, the Period transcript showed an opposite expression with the Timeless transcript in the second half of the free-running experiment, which was abnormal for the circadian clock regulated by the TTFL mechanism. Since a robust endogenous clock is required to be free running with a self-sustained period for maintaining the rhythmicity of tidal oscillations in *R. leurokolos*, we hypothesize that there is a circatidal clock with at least part of independent molecular components from the circadian clocks. Separate circadian and circatidal clocks were also reported in the intertidal isopod *Eurydice pulchra* [74] and mangrove cricket *Apteronemobius asahinai* [75]. Nevertheless, the core clock genes still encode functional proteins with conserved domains (electronic supplementary material, figure S12*e*), and we cannot exclude the possibility that some core circadian clock genes are necessary for tidal rhythms [76, 77], even though they lose free-running transcription rhythmicity. These circadian clock genes may also be useful during the dispersal of planktotrophic larvae, which may take place in the photic zone [78].

**Figure 4.**
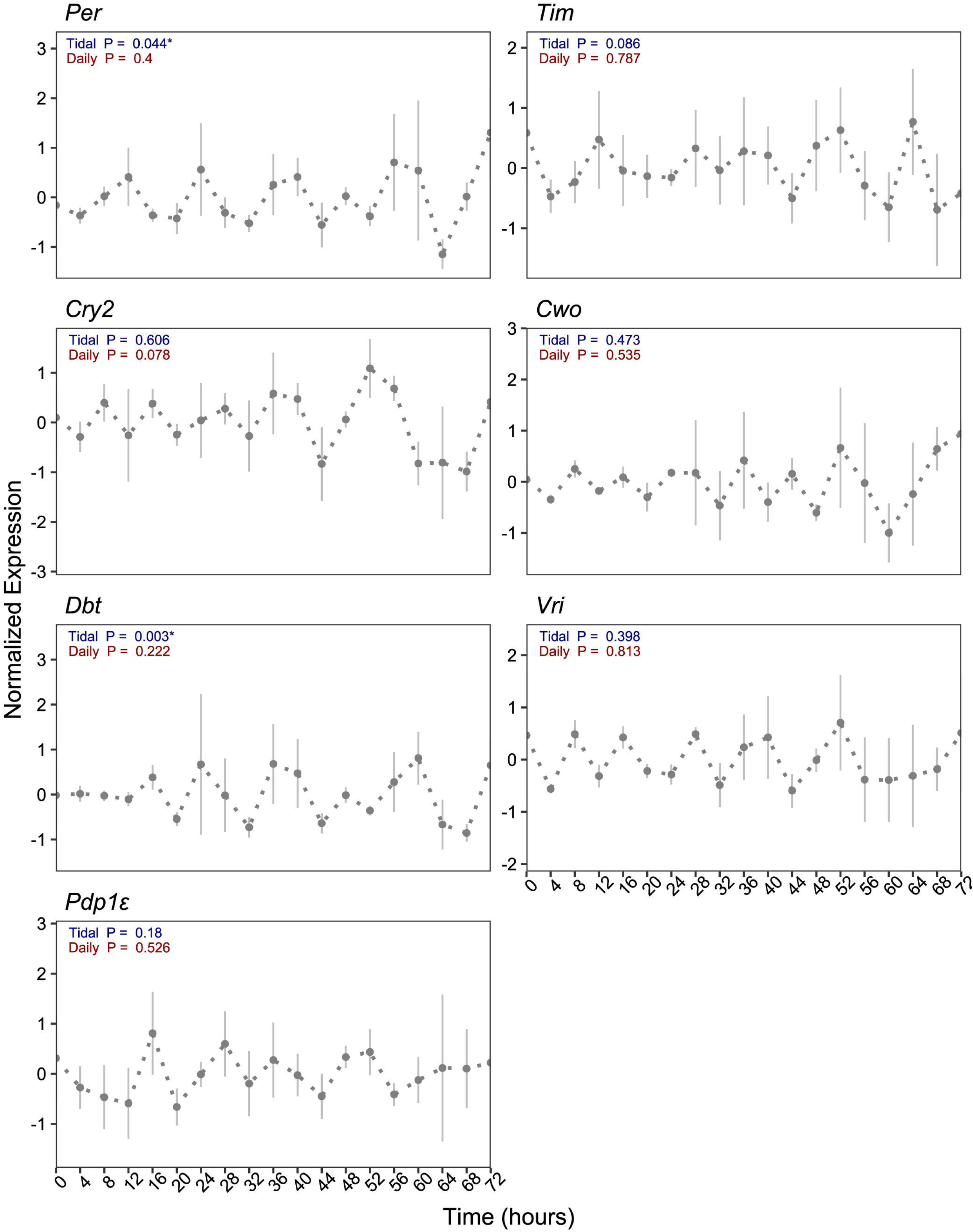
Expression profiles of *Rimicaris leurokolos* homologs of candidate circadian clock genes. Data points are displayed as mean values ± SEM (n = 3 biological replicates per time point). Minimum *P*-values of results from tidal or daily detection of each transcript are presented in the plot. Significant levels of oscillations are indicated with stars: * FDR < 0.05.

To precisely synchronize with the tidal cycle, we consider *R. leurokolos* shrimps depend on an endogenous circatidal clock to integrate all potential periodic cues. In the microhabitat where *R. leurokolos* lives, crucial environmental factors such as temperature, water chemistry, and dissolved oxygen can vary irregularly and dramatically in a very short time, generating noisy signals. Therefore, although these environmental factors are crucial for the survival of hot vent shrimps, they are likely poor periodic cues. Hydrostatic pressure and tidal currents, however, are largely in line with the standard of a synchronization cue [79, 80]. In addition, they show significant coherence with temperature, water chemistry, and dissolved oxygen [81]. The closer the distance from hot vents, the stronger of modulation of tidal pressure on environmental factors [17]. Therefore, we propose hydrostatic pressure as the most likely synchronization cue for *R. leurokolos*. Over longer time scales the overall levels of venting activity is known to be modulated by the tidal cycle. As such, by synchronizing to pressure, it is possible for hot vent shrimps to predict major environmental fluctuations. In addition to pressure and currents, light emissions from vents [82], food sources [32] or even soundscape cues from vents [83] could be potentially useful periodic cues for *R. leurokolos*. Nevertheless, how exactly hot vent shrimps synchronize their endogenous rhythms with environmental cycles remains a question for the future.

### (f) Rhythmicity comparison of two vent species living in different microenvironments

To compare the tidal rhythms in the deep-sea hydrothermal vent organisms, we applied our transcriptome analysis and rhythm detection pipeline to the published *in situ* gene expression dataset of the Atlantic deep-sea vent mussel *B. azoricus* [22]. Despite the different geographic localities of the two species, *B. azoricus* and *R. leurokolos* both show similar biological rhythms predominated by a tidal pattern that is related to the responses to environmental stresses such as DNA damage, and thermal, immune, and oxidative stress, suggesting a generally temporal adaptation to the harsh deep-sea hydrothermal environments.

Compared to *R. leurokolos*, however, the proportion of tidal transcripts in the transcriptome was lower in *B. azoricus* regardless of the detection algorithm used (electronic supplementary material, figure S13*a* and S13*b*). Indeed, considering that rhythmic gene expression was organ-specific [5], the result was disputable as we detected overall rhythmicity within mixed tissues of *R. leurokolos*, whereas the study on *B. azoricus* only worked on gills. We further examined and compared the strength of tidal rhythms in the two species by calculating the rhythmicity index (RI) using the autocorrelation analysis, and the results suggested the general rhythmicity of tidal gene expression was greater in *R. leurokolos* (electronic supplementary material, figure S13*c*). This is unexpected since it should be easier for *B. azoricus*, living in a relatively distant area from the vent orifice with stable diffuse flow, to detect environmental factors influenced by tidal cycles. A possible explanation is that *R. leurokolos* needs to depend on a strong pacemaker to integrate all the periodic cues for predicting environmental fluctuations and preparing for transcriptional regulation, whereas *B. azoricus* could schedule the transcription by either relying on an endogenous clock or responding directly to the tidal-modulated environmental cues [22]. Why *R. leurokolos* inhabiting a more dynamic environment shows more robust rhythmicity compared to *B. azoricus* is an intriguing topic warranting future studies, although this could also be related to intrinsic differences between mussels and shrimps, as well as the influence of chemosymbiosis on the host [84].

### (g) The potential evolutionary conservation between marine tidal rhythms and terrestrial 12-hour rhythms

We show that the deep-sea vent shrimp *R. leurokolos* exhibits robust endogenous rhythms related to tidal effects instead of daily cycles, making it a useful model for testing the hypothesis that marine tidal rhythms and 12-hour rhythms of terrestrial animals share conserved features. To investigate this, we compared the tidal gene expression of *R. leurokolos* with published ∼12-hour rhythms in the *Bmal1* knockout mice liver [28] and *Drosophila* S2 cells [29], two systems without canonical circadian clock gene expressions. A total of 406 single-copy orthologous genes were identified as showing ∼12-hour rhythms in these three species. The vent shrimp *Rimicaris* shared more genes with the fruit fly compared with the mouse (54 genes vs 36 genes), suggesting a phylogenetic relationship. We also found 21 genes with a period of ∼12-hour rhythms in all three species (figure 5*a*). These genes were enriched in the processes of DNA replication, regulation of transcription, ribosome biogenesis, protein folding, and trans-Golgi network, which are also highly enriched in the ∼12-hour rhythms of human beings [85] (figure 5*b*). The heat shock protein 70 cochaperone *Dnaja1*, which was also reported in 12-hour rhythms of mammals [85] and tidal rhythms of the sea anemone *Aiptasia diaphana* [84], also showed a robust free-running tidal rhythm in *R. leurokolos*. These results support the evolutionary conservation of ∼12-hour gene expression related to the processes of central dogma information flow in both terrestrial and marine animals across a broad range of phyla, indicating the ∼12-hour rhythms in animals overall likely initially evolved for adjusting to the tidal cycle.

**Figure 5.**
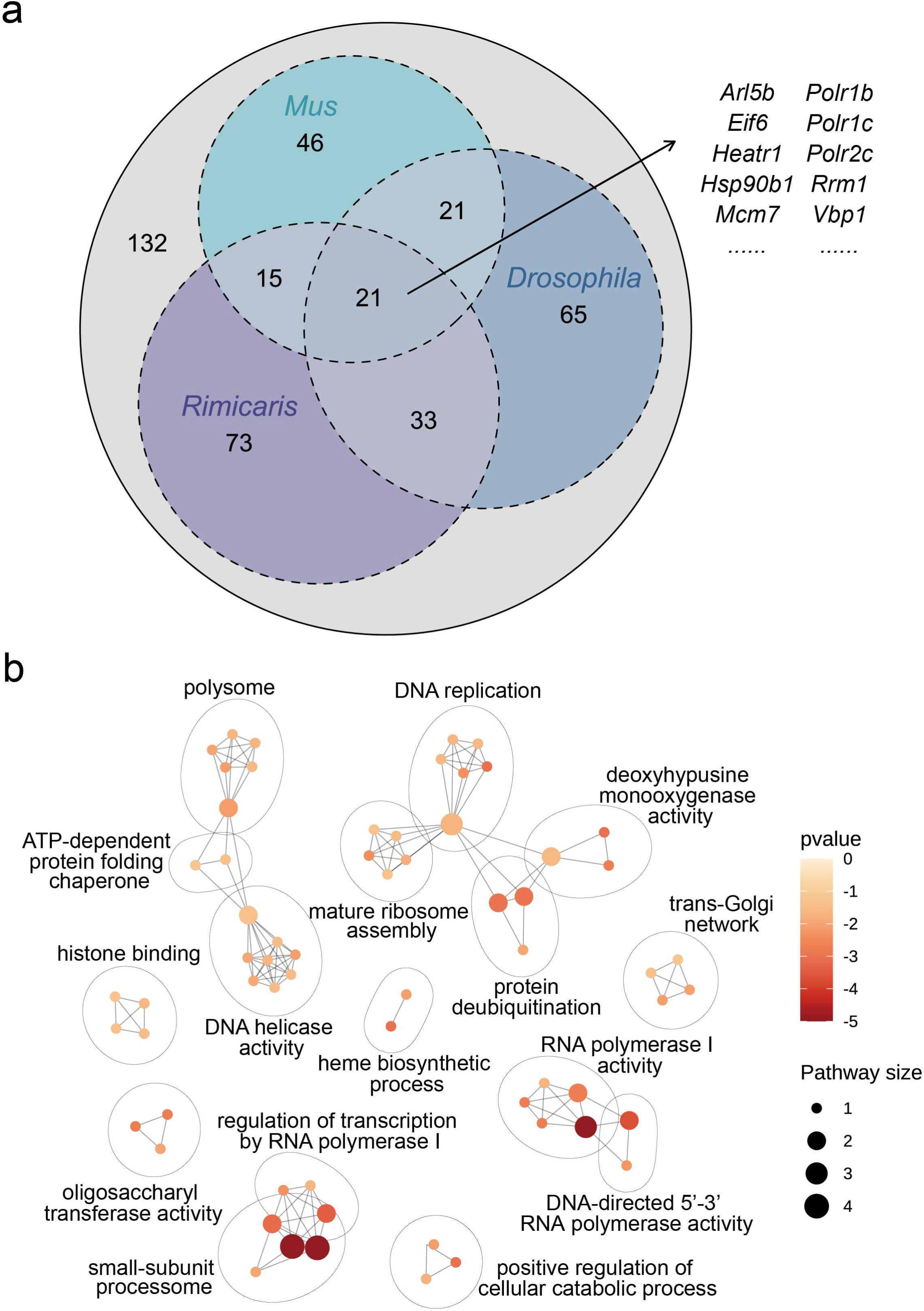
Homologous genes with a period of 12-hour rhythms between *Rimicaris leurokolos*, *Mus musculus*, and *Drosophila melanogaster*. (*a*) Venn diagram showing the comparison of ∼12-hour rhythms of single-copy orthologous genes between three species. (*b*) Pathway enrichment network showing biological functions of 21 conserved ∼12-hour genes in three species.

## 4. Conclusion

In this study, our analyses provide evidence for endogenous biological rhythms in the deep-sea hydrothermal vent shrimps *R. leurokolos* through a 72-hour free-running experiment and constrain the possible zeitgeber for biological clocks ticking in the deep, shedding light on biological rhythms in the light-absent biosphere (figure 6). We propose that rhythms with a period of 24 hours driven by the conventional circadian clock largely decrease or even disappear in adult deep-sea hydrothermal vent fauna. Instead, rhythmic gene expression coordinated with approximately 12-hour tidal cycles, which are likely controlled by an independent circatidal clock, evolved as a temporal adaptation to the environmental fluctuations in hydrothermal ecosystems. Independent of solar energy but being influenced by tidal cycles, organisms in the deep-sea hydrothermal vent are likely powerful models for chronobiologists to decipher the evolution and molecular mechanisms of circatidal clocks and ∼12-hour rhythms, important across a wide range of living organisms.

**Figure 6.**
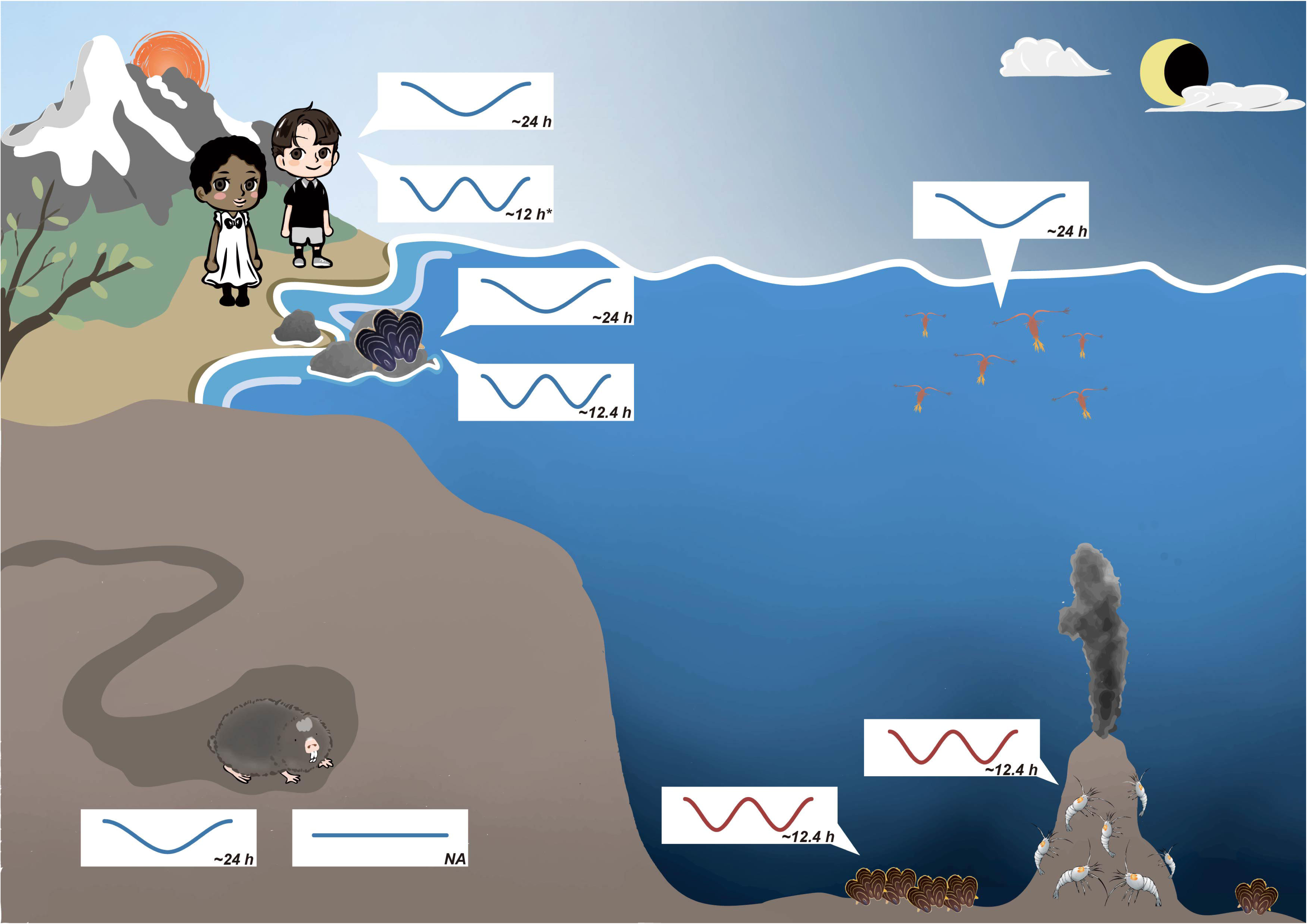
A new perspective of biological rhythms in the dark biosphere. Terrestrial and shallow-water organisms largely rely on circadian clocks for anticipating periodic environmental changes such as the day-night transition. Terrestrial organisms such as human beings exhibit robust ∼24-hour rhythms driven by the circadian clock [2], and potential ∼12-hour ultradian rhythms (with asterisk) that are assumed to be homologous with circatidal rhythms [86], though the biological functions and synchronization cues are still unclear. The intertidal mussel *Mytilus californianus* shows both circatidal and circadian rhythms, while the latter is the dominant one [51]. The copepod *Calanus finmarchicus* also shows behavioral rhythms in diel vertical migration (DVM) [87]. However, a different scenario can be found in the dark biosphere. Living away from direct sunlight, subterranean mole rats scarcely depend on circadian rhythms for survival, and the endogenous circadian system is understood to be in the process of regression, even though still present [88]. In this work, we found that in addition to the vent mussel *B. azoricus* [22], the vent shrimp *Rimicaris leurokolos* shows strong endogenous 12-hour rhythms in the gene expression, likely reflecting important effects of tidal cycles on deep-sea hot vent organisms.

## Data accessibility

Raw RNA-Seq data are available on NCBI as BioProject PRJNA993859. The *de novo* reference transcriptome of *Rimicaris leurokolos* is available in figshare (https://doi.org/10.6084/m9.figshare.23667327). All original codes used in this work are available at GitHub (https://github.com/belphe/Shinkaicaris).

## Author’s contributions

Conceptualization, J.S., T.Y., N.M., and C.C.; Methodology, T.Y., N.M., C.C., and H.Y.Z.; Formal Analysis, H.Y.Z. and J.S.; Investigation, T.Y., C.C.; Writing – Original Draft, H.Y.Z.; Writing – Review & Editing, J.S., C.C.; Visualization, H.Y.Z. and Q.Q.J.; Supervision, J.S. and T.Y.; Project Administration, J.S., P-Y.Q. and T.Y.; Funding Acquisition, J.S. and P-Y.Q.

## Competing interests

The authors declare that they have no competing interests.

## Funding

This work was supported by: Science and Technology Innovation Project of Laoshan Laboratory (LSKJ202203104); Fundamental Research Funds for the Central Universities (202172002 and 202241002); Young Taishan Scholars Program of Shandong Province (tsqn202103036); Natural Science Foundation of Shandong Province (ZR2023JQ014); Principal investigator projects of the Southern Marine Science and Engineering Guangdong Laboratory (Guangzhou) (2021HJ01); Major Basic and Applied Research Projects of Guangdong Province (2019B030302004-04); Southern Marine Science and Engineering Guangdong Laboratory (Guangzhou) (SMSEGL20SC01); Hong Kong Special Administrative Region government (16101822, C2013-22GF)

## Acknowledgements

We are grateful to the ship captain and crews of R/V *KAIYO* as well as pilots and the technical team of ROV *Hyper-Dolphin* for their great efforts during sample collection during cruise KY14-01 led by Ken Takai (JAMSTEC). Hiroyuki Yamamoto, Shinsuke Kawagucci, and Hiromi K. Watanabe (JAMSTEC) are thanked for providing raw data of the temperature probe recordings published in a previous study [31]. We also thank Bokai Zhu (University of Pittsburgh) for his insightful advice on an earlier version of this manuscript.

## Notes

### Competing Interest Statement

The authors have declared no competing interest.

### Summary of Updates

An in-depth comparison of the raw gene expression data and the "detrended" gene expression data; All the Figures revised; The whole manuscript further revised; Supplementary filed updated.

