## Supplementary Material for "Circatidal control of gene expression in the deep-sea hot vent shrimp *Rimicaris leurokolos*"

**The Electronic Supplementary Materials include:**

• Supplementary Figures S1 to S13

• Supplementary Tables S1

**Supplementary Figures**

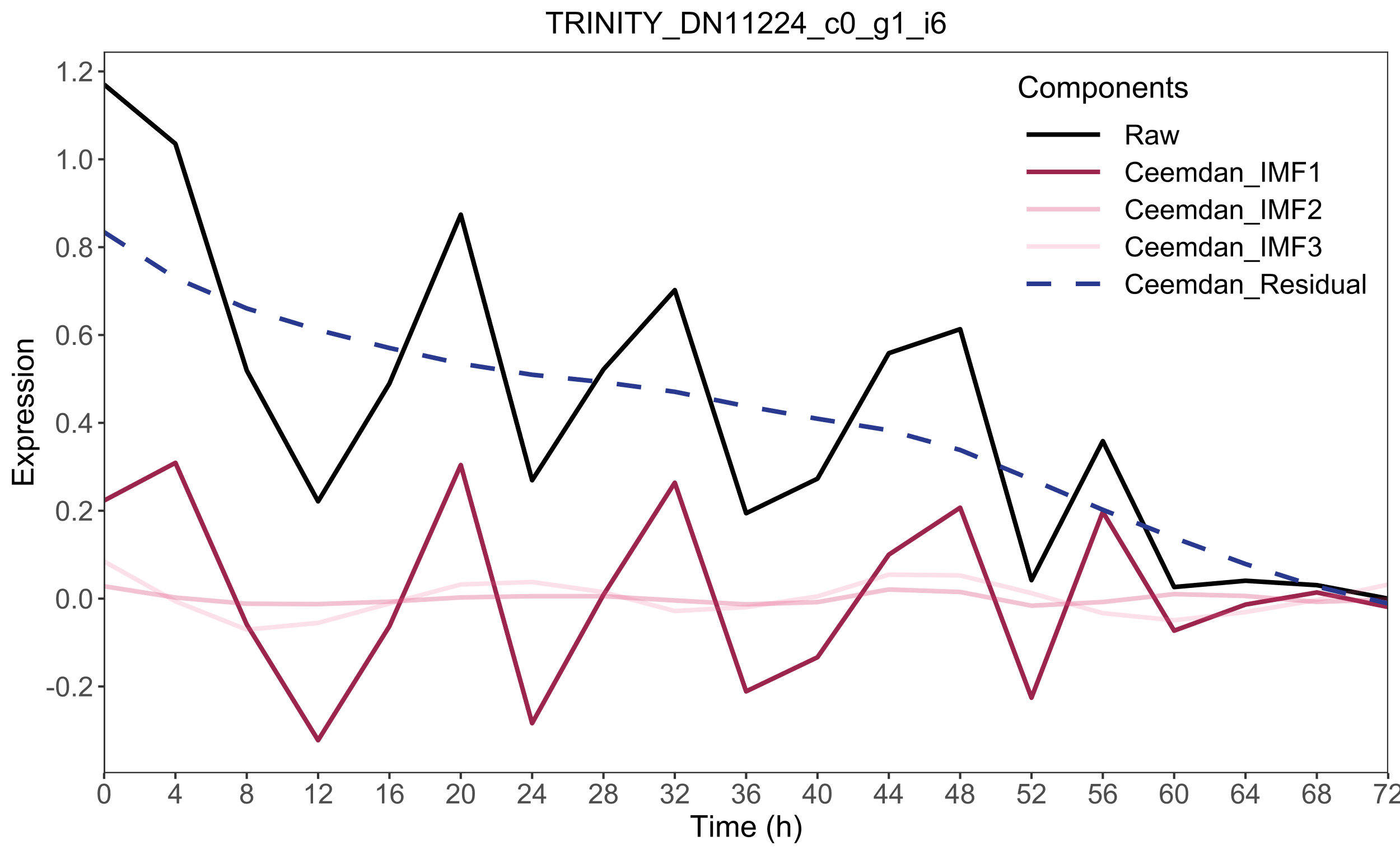

**Supplementary Figure 1.** An example about the detrend analysis based on CEEMDAN. The raw gene expression is decomposed into 3 IMFs that may contain potential periodic patterns and a residual that represents the trend. For this transcript, the component “Ceemdan_IMF1” is considered as the detrended gene expression under criterion.

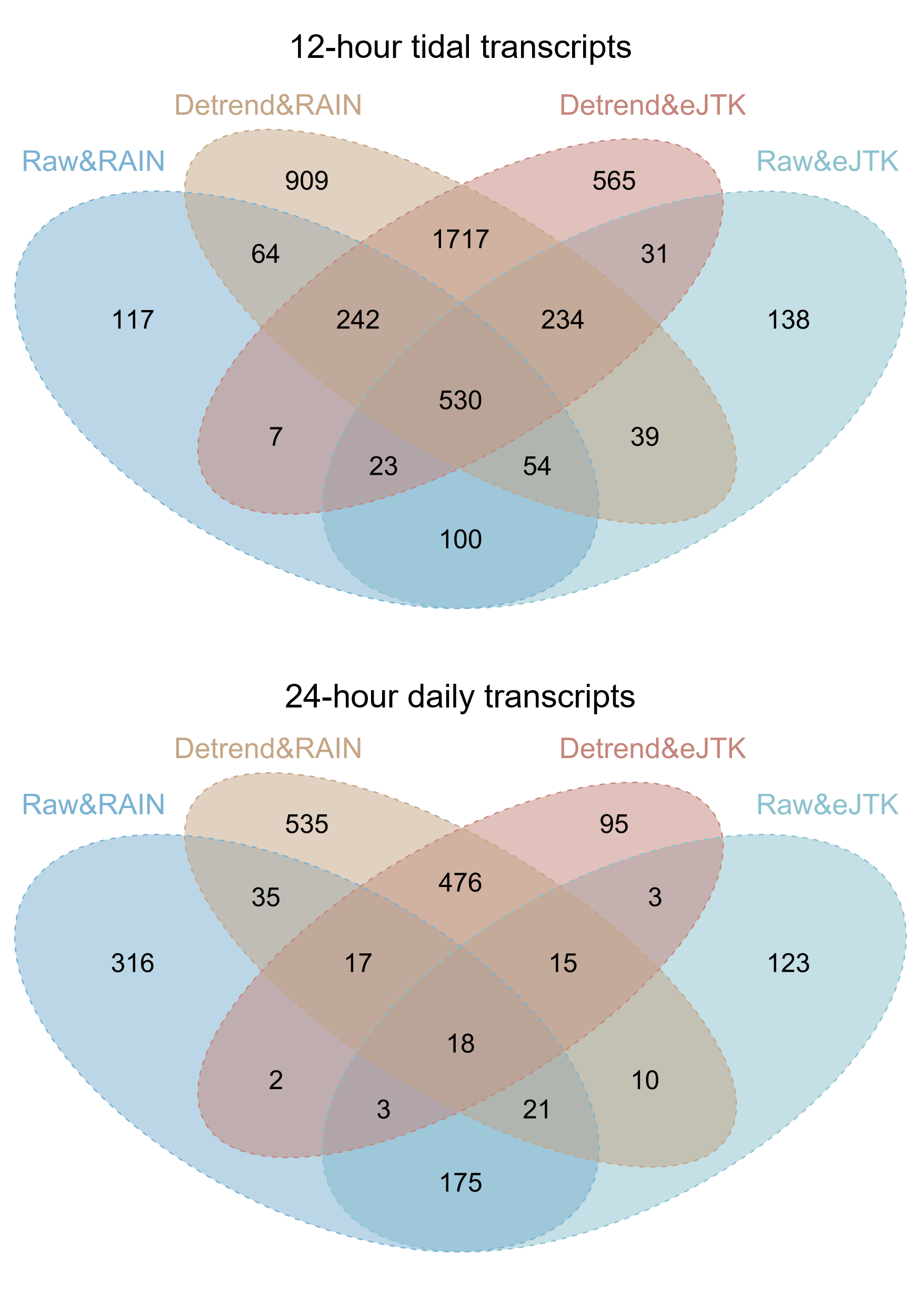

**Supplementary Figure 2.** Venn diagrams show the comparison of the number of 12-hour tidal transcripts and 24-hour daily transcripts that are identified using gene expression data before and after detrending.

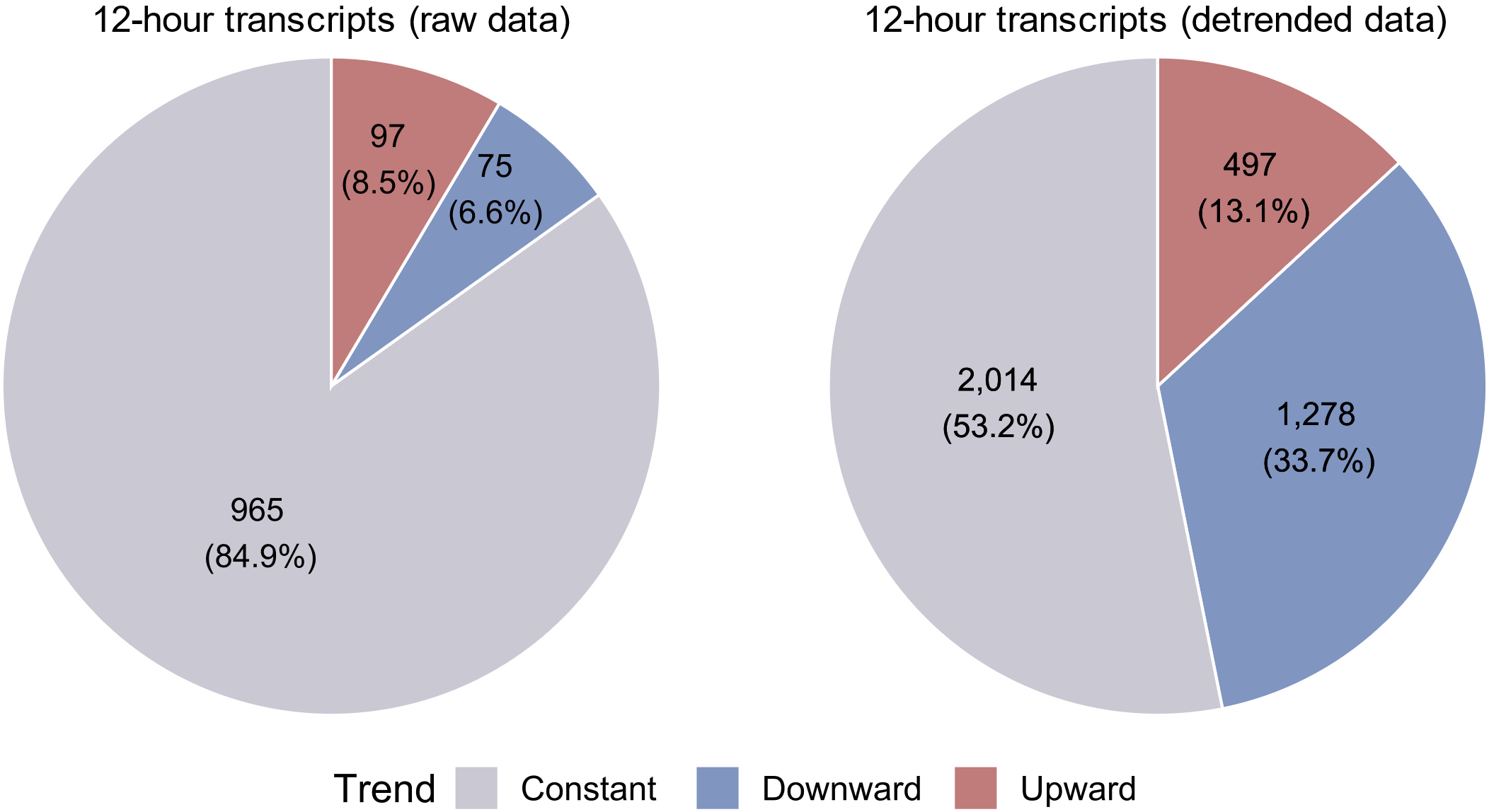

**Supplementary Figure 3.** Pie charts showing the number and proportion of 3 trend types in the tidal transcripts identified by RAIN using raw and detrended gene expression data.

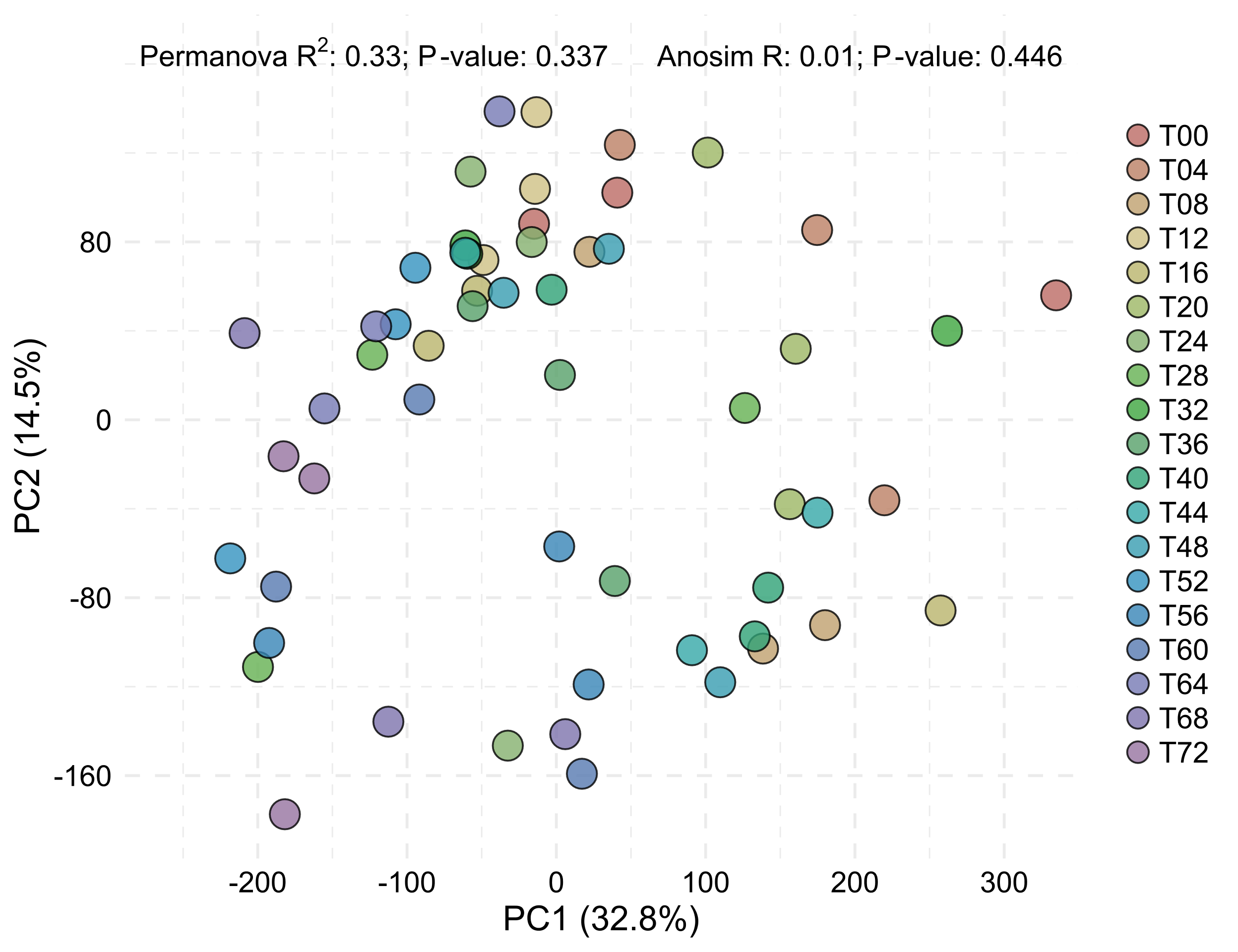

**Supplementary Figure 4.** Principal components analysis (PCA) plot showing the detrended transcriptomic data from the free-running experiment samples (n = 57). Significance of inter-group differences was examined by the PERMANOVA (*P*-value = 0.337) and ANOSIM (*P*-value = 0.446) analysis both on Bray-Curtis dissimilarities.

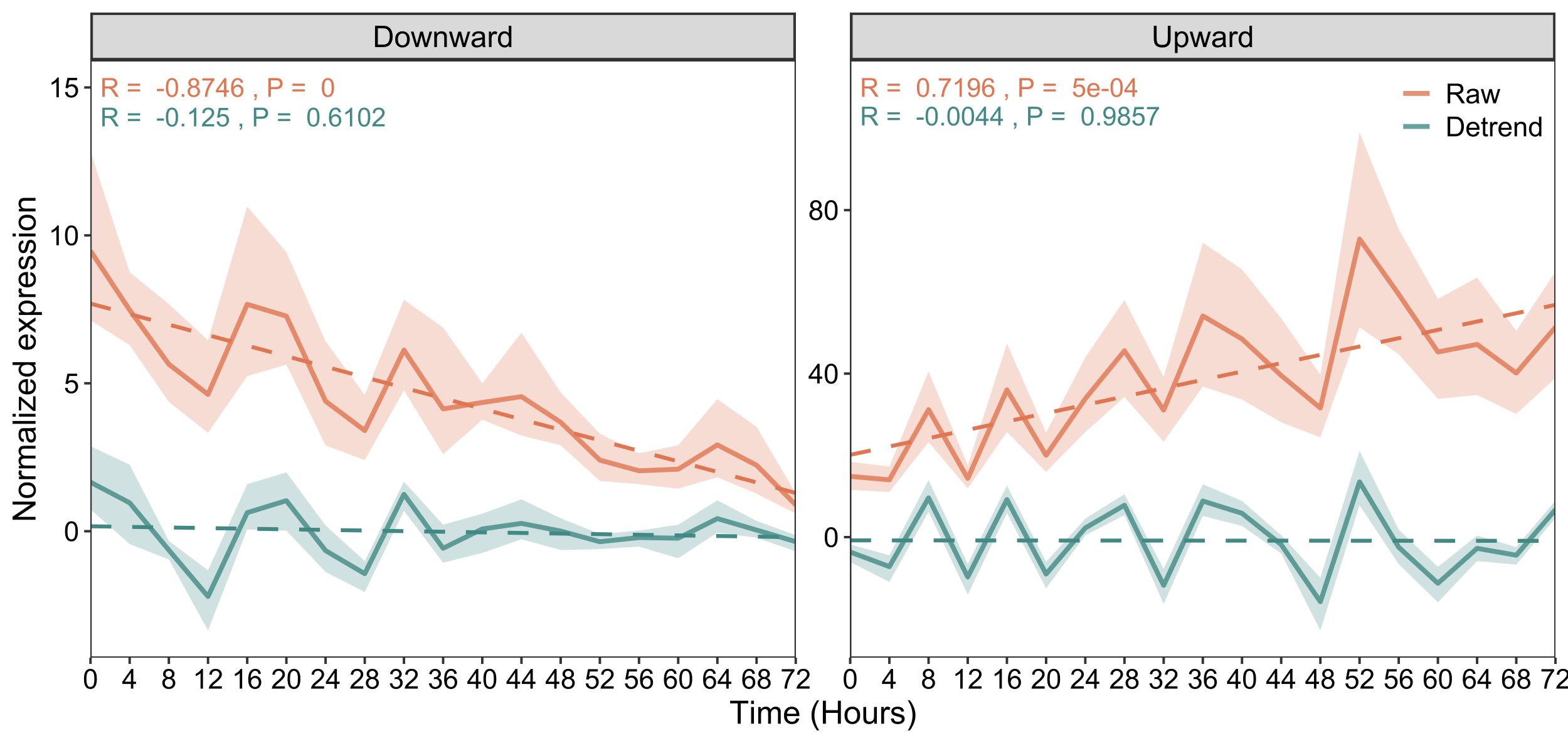

**Supplementary Figure 5.** Expression plots of genesets with upward or downward trends before and after detrending. Significance of correlation is tested with the Pearson correlation coefficient.

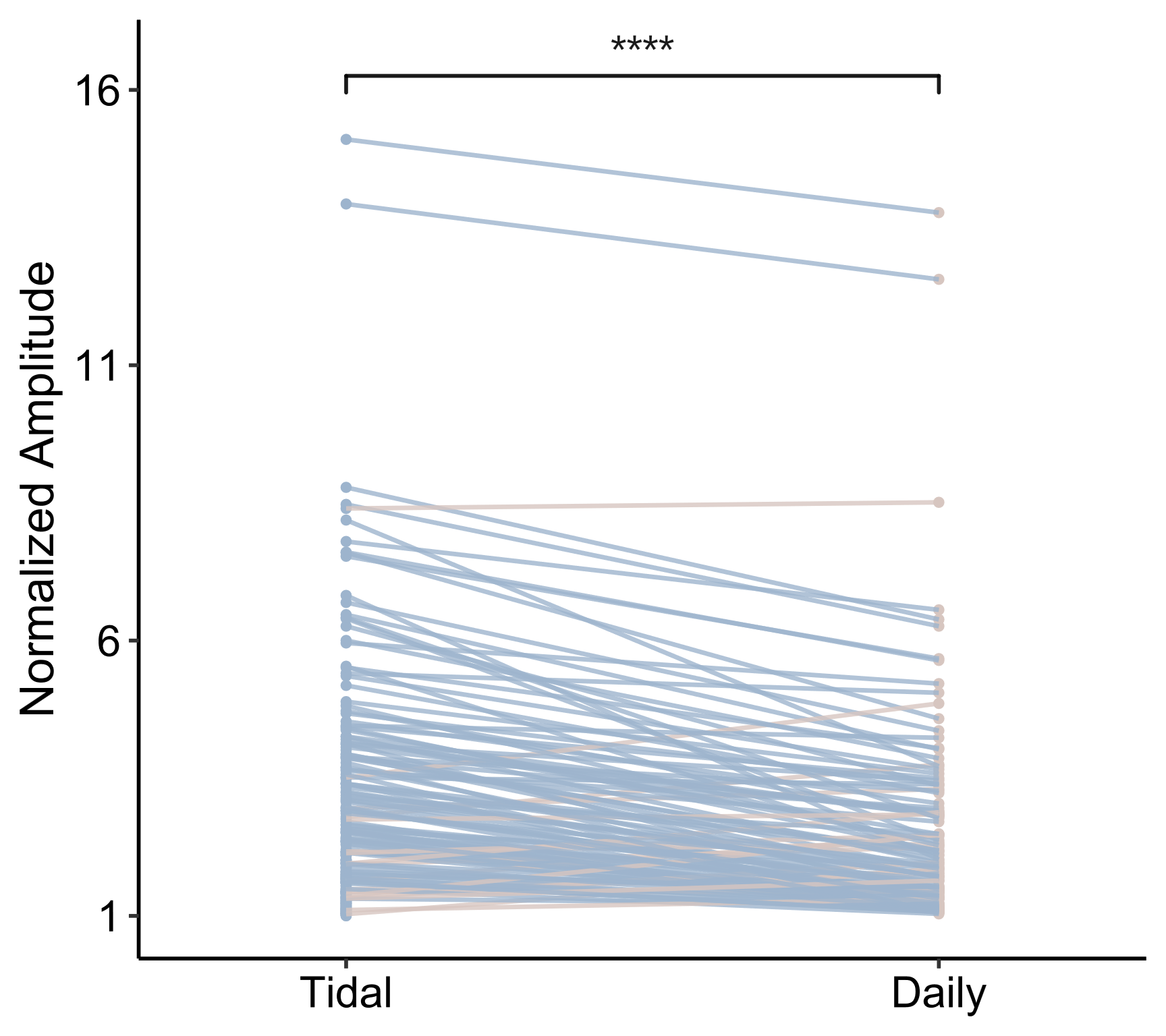

**Supplementary Figure 6.** Paired t-test plot of amplitude for tidal and daily components of superimposed cycling transcripts (identified by the eigenvalue/pencil method) that are comprised of multiple components composed of tidal and daily oscillations. The stars indicated the level of significance: **** *P*-value < 0.0001.

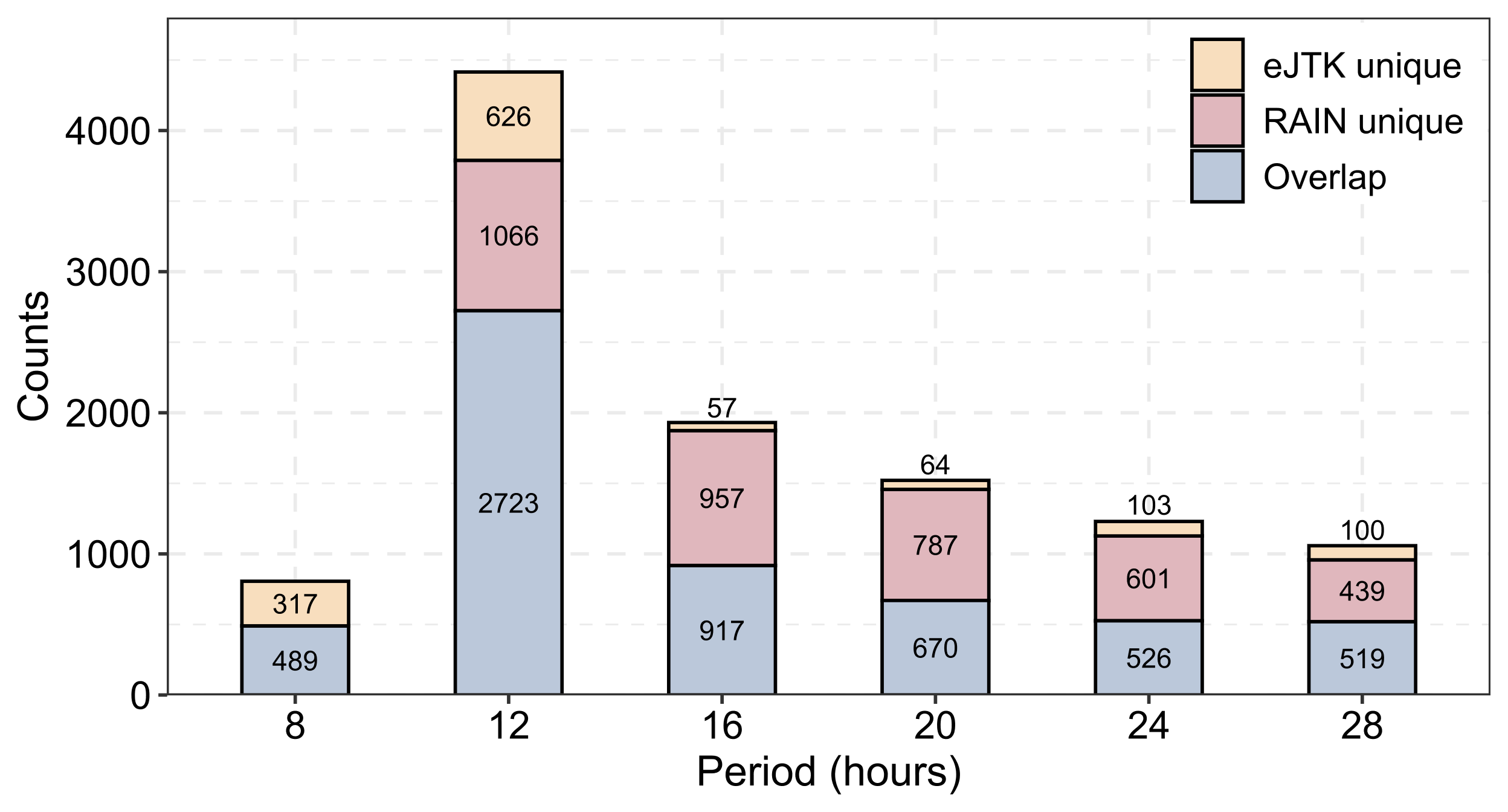

**Supplementary Figure 7.** Bar chart showing the distribution of periods of rhythmic transcripts detected by RAIN and eJTK (FDR < 0.05), related to figure 2.

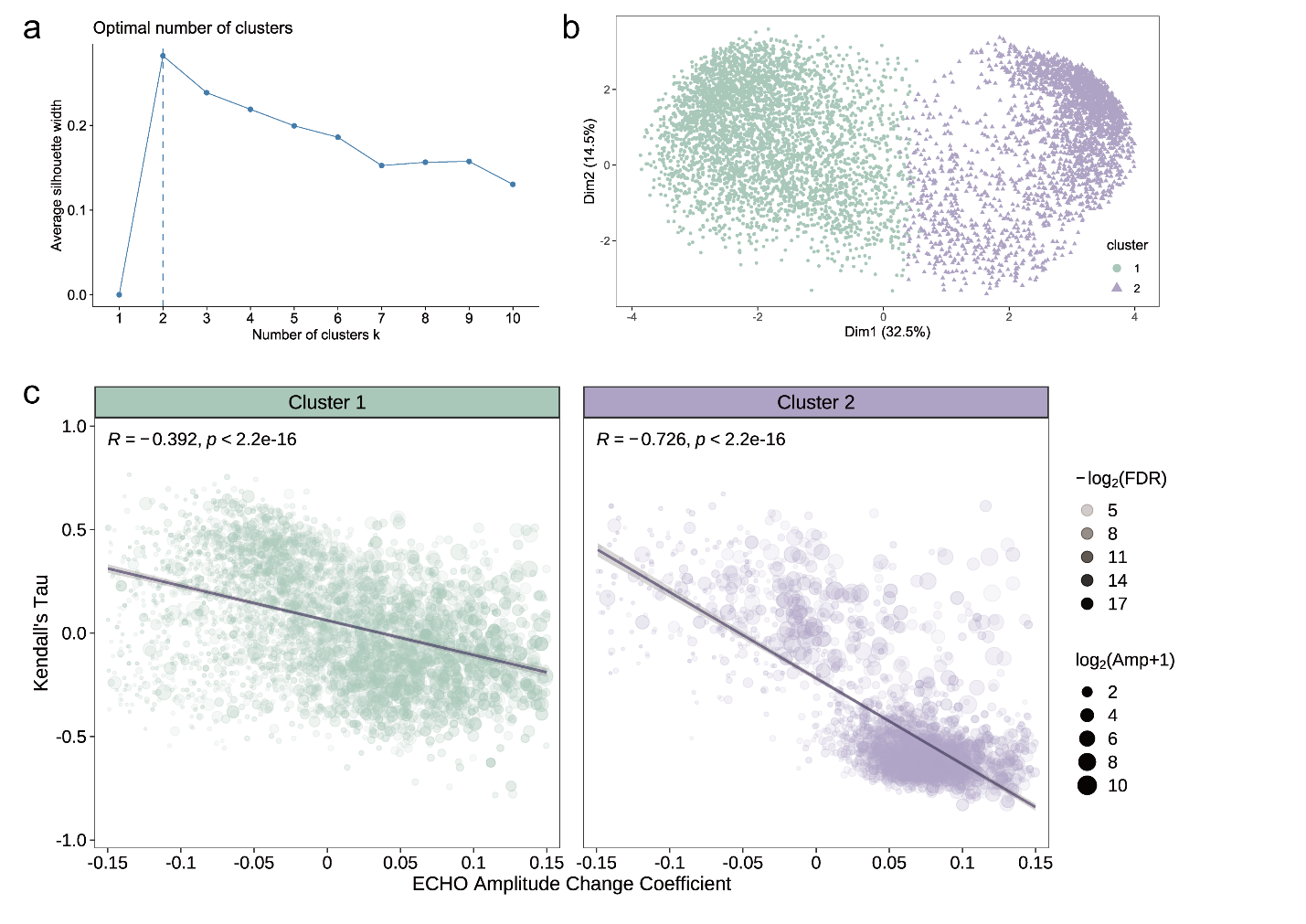

**Supplementary Figure 8.** Two antiphased clusters of tidally oscillated transcripts. (*a,b*) Kmeans clustering based on detrended gene expression of tidal transcripts. (*a*) Silhouette validation measure shows kmeans method with (*b*) two clusters is the most optimal one. (*c*) Scatter diagram showing the correlation between the ECHO amplitude change coefficient and Kendall’s tau. Data is fitted with a linear regression. Significance of correlation is tested with the Pearson correlation coefficient.

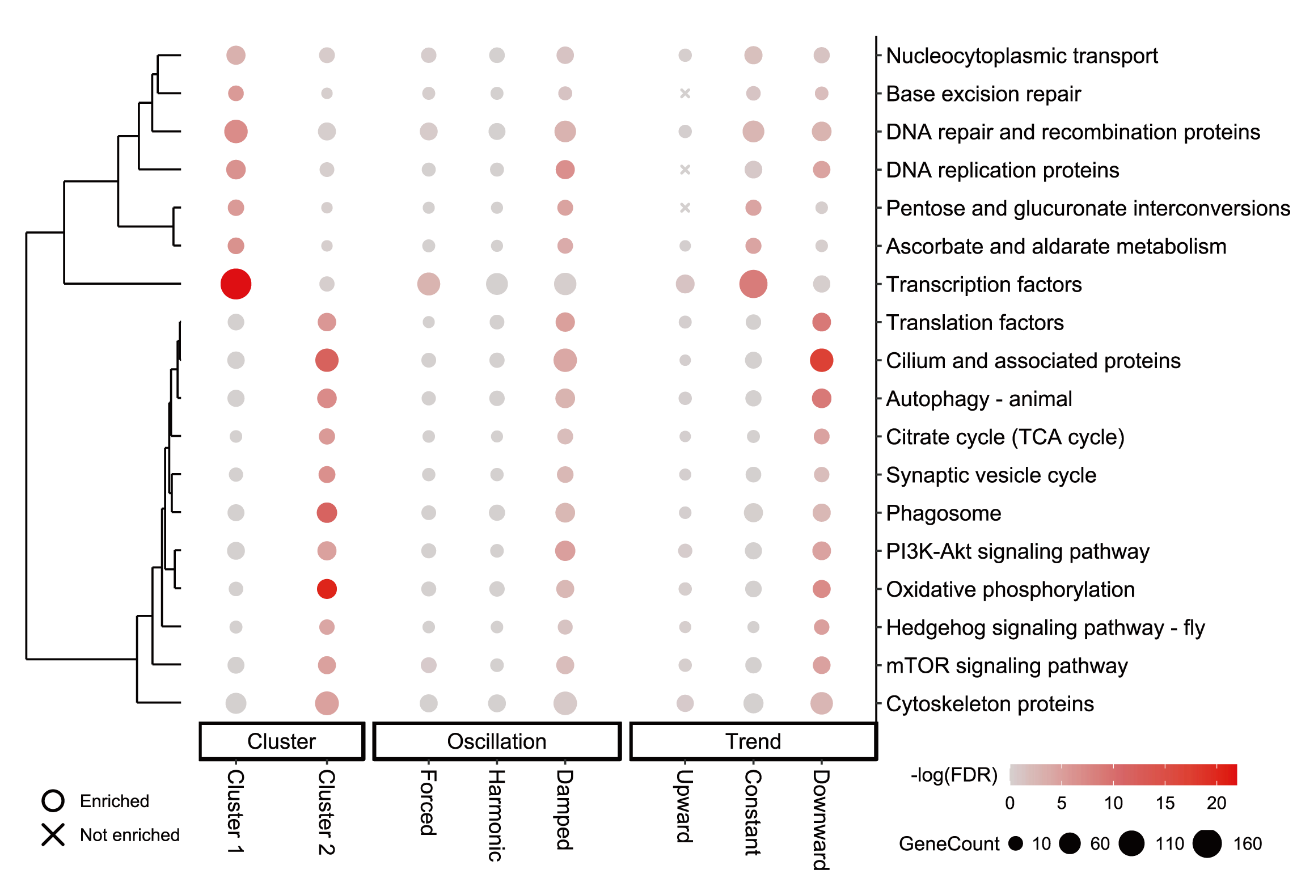

**Supplementary Figure 9.** Bubble diagram showing significantly enriched KEGG terms, related to figure 3. Tidal subgroups are categorized by three classification schemes (from left to right: cluster, oscillation type, trend). Cross symbols represent a lack of enriched terms in the corresponding subgroup.

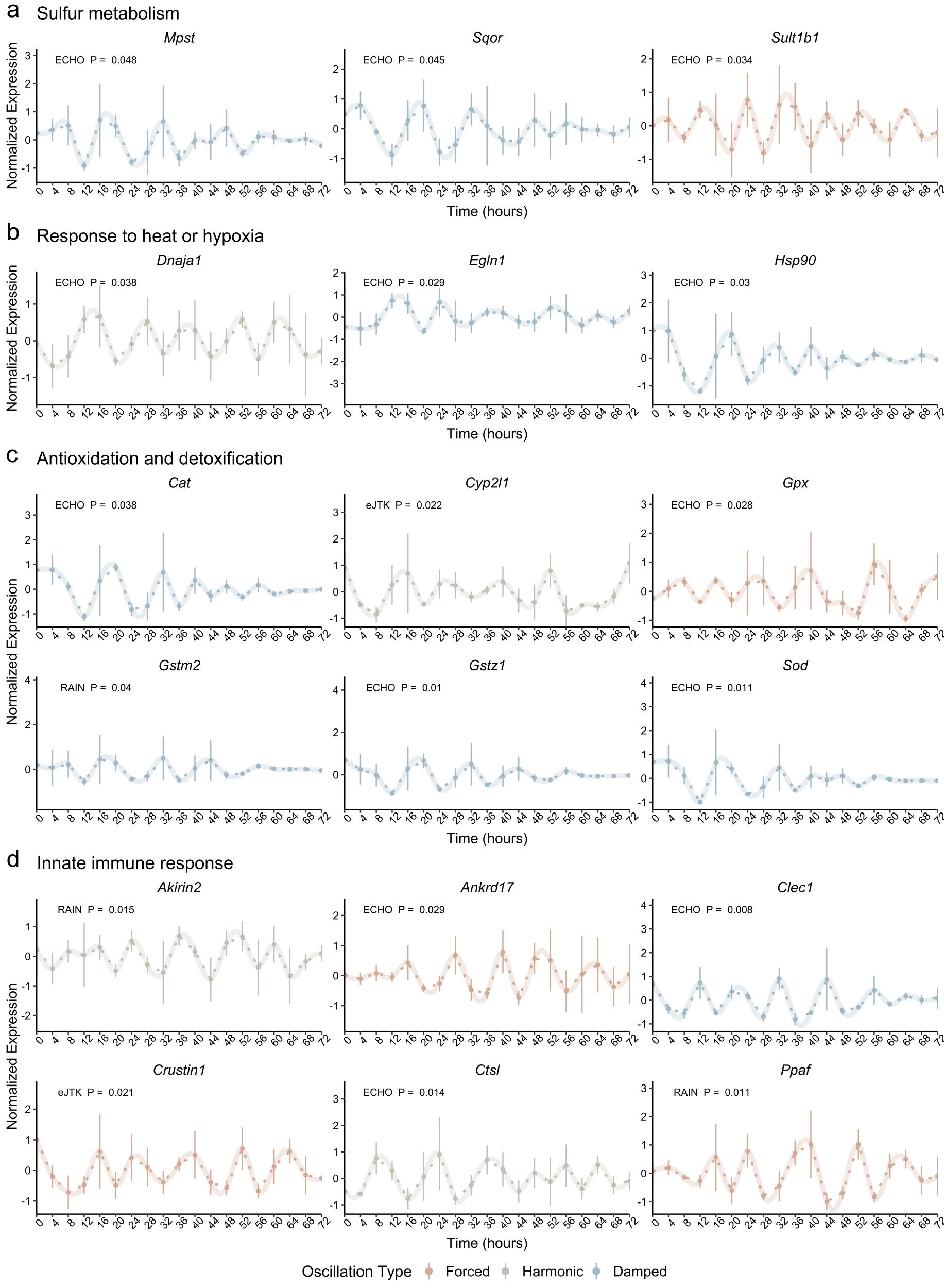

**Supplementary Figure 10**. Expression profiles of examples of tidal transcripts of interest. The listed tidal genes are potentially involved in various cellular stress response processes, including (a) sulfur metabolism, (b) response to heat or hypoxia, (c) antioxidation and detoxification, and (d) innate immune response. Data points are displayed as mean values ± SEM (n = 3 biological replicates per time point). Normalized expression curves are fitted via the generalized additive model (GAM). Minimum *P*-values of each tidal transcript identified by RAIN, eJTK, and ECHO are presented in the plot.

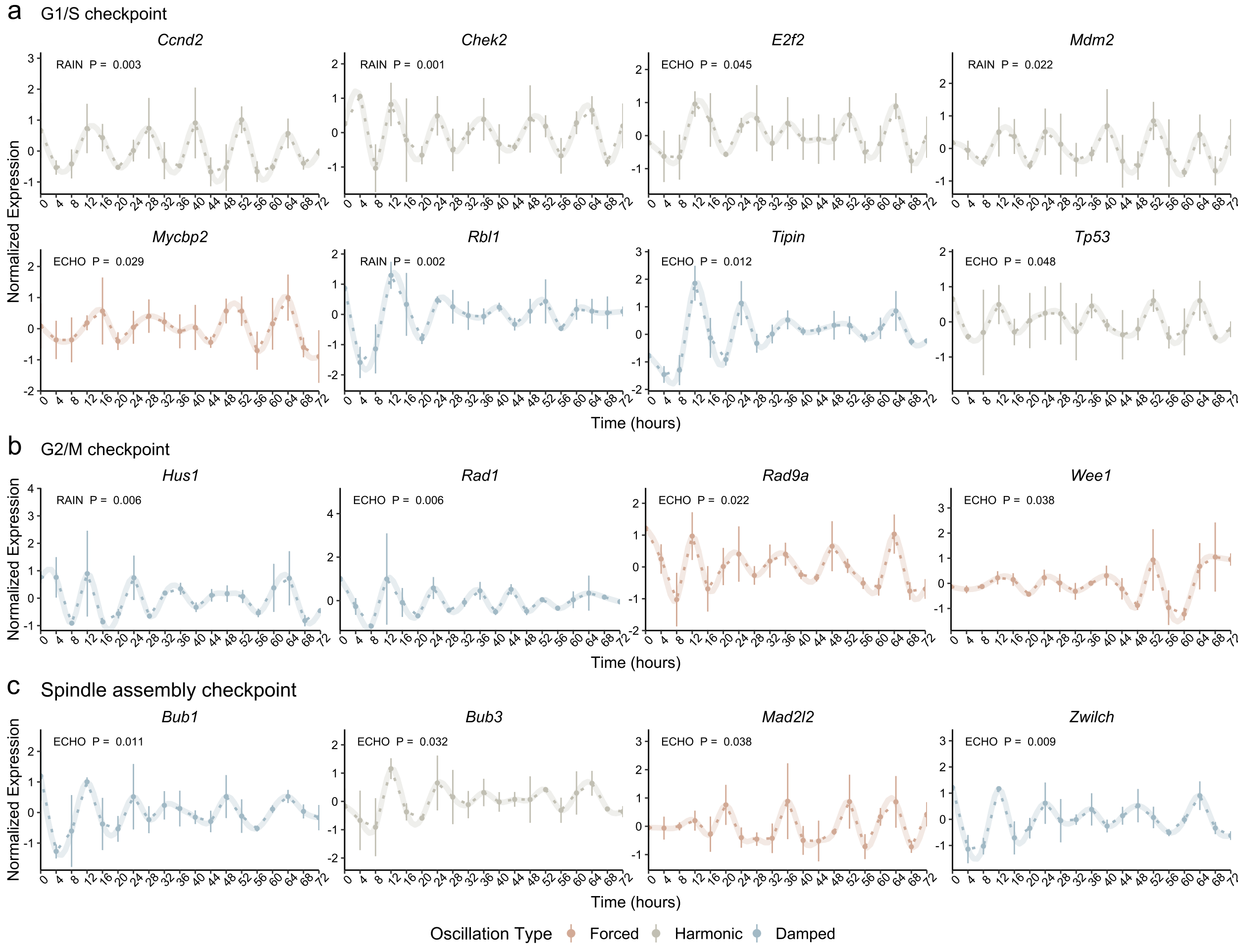

**Supplementary Figure 11.** Expression profiles of examples of tidal transcripts related to the cell cycle. The listed tidal genes potentially play crucial roles in cell cycle checkpoints, including (*a*) G1/S checkpoint, (*b*) G2/M checkpoint, and (*c*) spindle assembly checkpoint (SAC). Data points are displayed as mean values ± SEM (n = 3 biological replicates per time point). Normalized expression curves are fitted via the generalized additive model (GAM). Minimum *P*-values of each tidal transcript identified by RAIN, eJTK, and ECHO are presented in the plot.

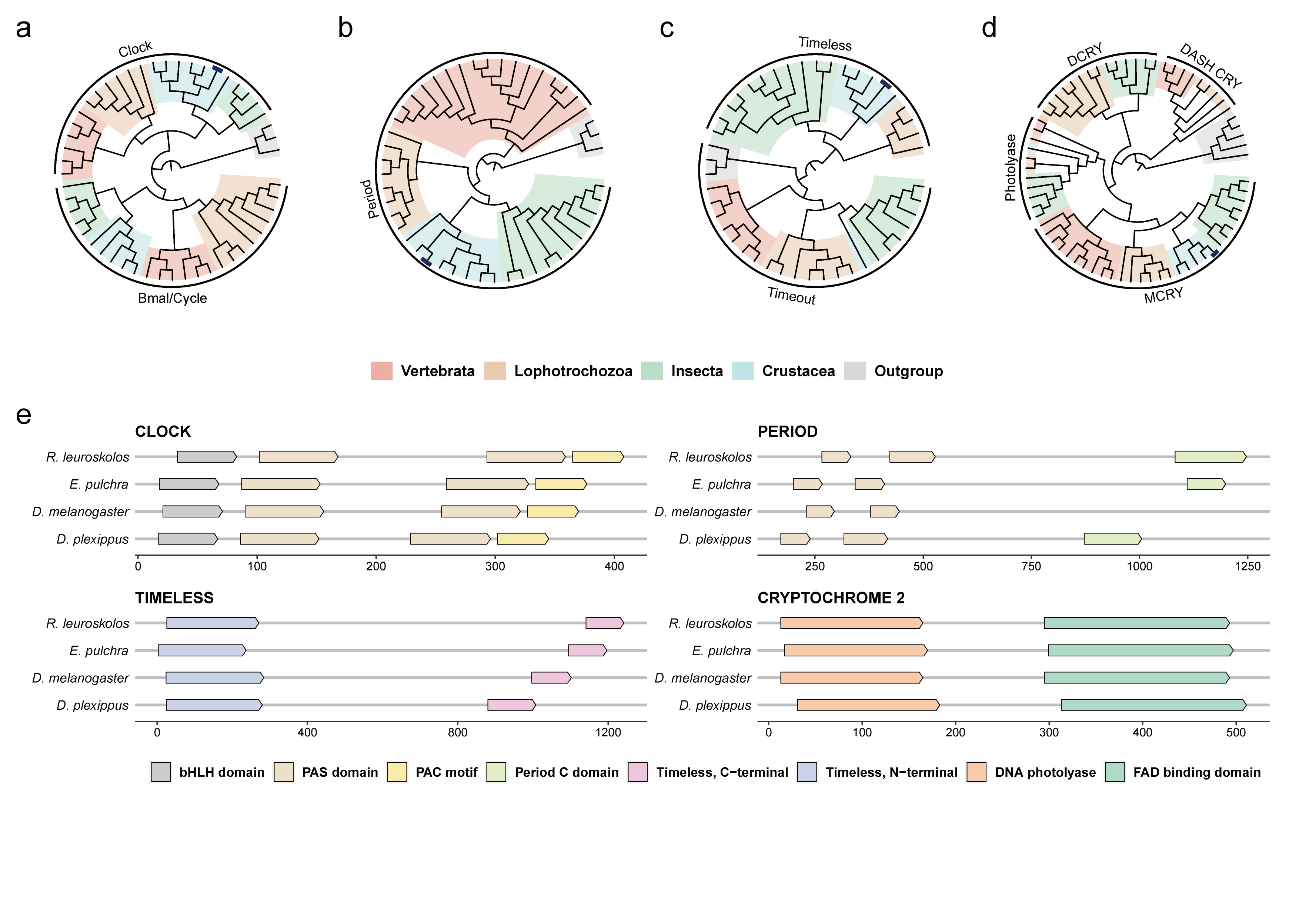

**Supplementary Figure 12.** Putative core circadian clock genes identified in *R. leurokolos*. (*a-d*) Simplified Maximum-likelihood phylogenetic relationships of (*a*) Clock and Bmal/Cycle protein family, (***b***) Period protein family, (***c***) Timeless and Timeout protein family, and (***d***) Photolyase/Cryptochrome protein family. *R. leurokolos* proteins are highlighted in dark blue. (***e***) Schematic presentation of functional domains of the *R. leurokolos* core clock proteins. Domains structure is compared to *Eurydice pulchra*, *Drosophila melanogaster*, and *Danaus plexippus*. Grey bars indicate amino acidic length sequence. Specific protein domains identified by the SMART analysis are colored.

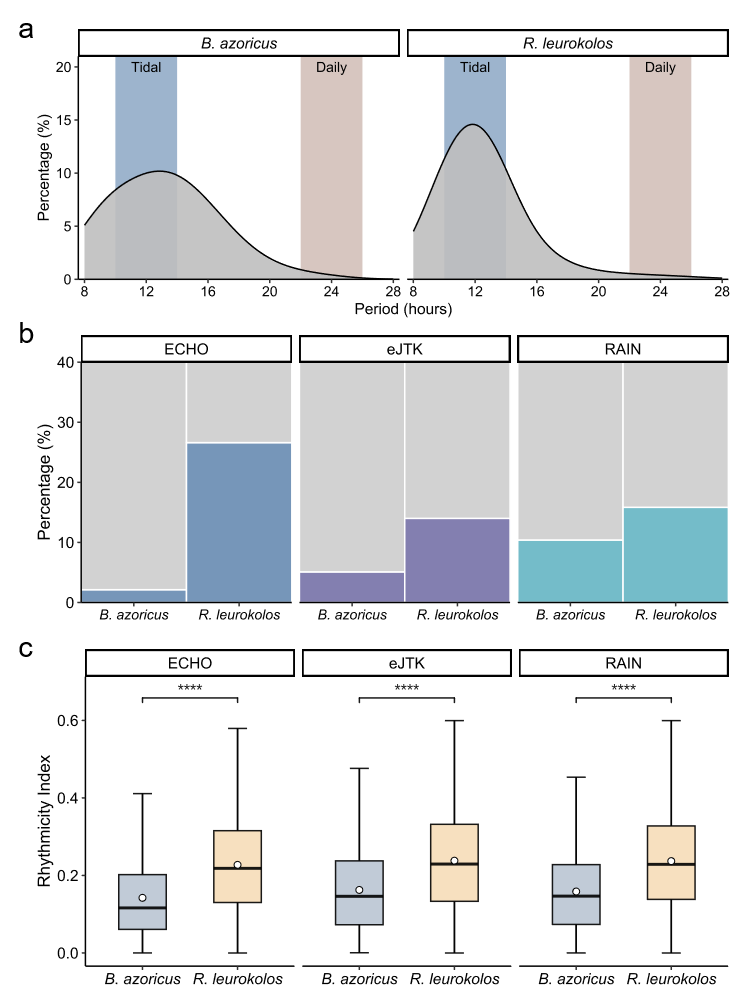

**Supplementary Figure 13.** Tidal transcriptional rhythmicity comparison of the vent shrimp *Rimicaris leurokolos* and mussel *Bathymodiolus azoricus*. (*a,b*) Comparison of the proportion of rhythmic transcripts in the temporal transcriptome of *R. leurokolos* and *B. azoricus*, which are identified by (*a*) the eigenvalue/pencil method and (*b*) RAIN, eJTK, and ECHO. Rhythmic transcripts are Oscillations with a period range of 10-14 hours and 22-26 hours are regarded as tidal and daily cycling, respectively. (*c*) Box graph showing the comparison between rhythmicity index (RI) for the tidal transcripts of *B. azoricus* and *R. leurokolos*. Peaks in the autocorrelation corresponding to a lag of 12 hours (for *R. leurokolos*) and 12.4 hours (for *B. azoricus*) are used to calculate the rhythmicity index. The stars indicated the level of significance: **** *P*-value < 0.0001.

**Supplementary Tables**

**Supplementary Table 1** Comparison of statistical rhythm detection methods

| **Method** | **Statistical test type** | **Replicates** | **Asymmetric waveforms** | **Oscillation types** | **References** |
| --- | --- | --- | --- | --- | --- |
| Lomb-Scargle | Parametric | Yes | No | No | [1] |
| GeneCycle | Parametric | No | No | No | [2] |
| ARSER | Parametric | No | No | No | [3] |
| JTK_CYCLE | Non-parametric | Yes | No | No | [4] |
| RAIN | Non-parametric | Yes | Yes | No | [5] |
| eJTK | Non-parametric | Yes | Yes | No | [6] |
| ABSR | Parametric | No | No | No | [7] |
| BIO_CYCLE | Parametric | Yes | No | No | [8] |
| MetaCycle | Parametric | Yes | No | No | [9] |
| ECHO | Parametric | Yes | No | Yes | [10] |

Adapted from Mei et al. [11]
